# Integrating collecting systems in kidney organoids through fusion of distal nephron to ureteric bud

**DOI:** 10.1101/2024.09.19.613645

**Authors:** Min Shi, Brittney Crouse, Nambirajan Sundaram, Naomi Pode Shakked, Lioba Ester, Weitao Zhang, Vinothini Janakiram, Raphael Kopan, Michael A. Helmrath, Joseph V. Bonventre, Kyle W. McCracken

**Affiliations:** Division of Nephrology and Hypertension, Department of Pediatrics, Cincinnati Children’s Hospital Medical Center, Cincinnati, OH, USA; Center for Stem Cell and Organoid Medicine, Cincinnati Children’s Hospital Medical Center, Cincinnati, OH, USA; Department of Surgery, Cincinnati Children’s Hospital Medical Center, Cincinnati, OH, USA; Division of Developmental Biology, Department of Pediatrics, Cincinnati Children’s Hospital Medical Center, Cincinnati, OH, USA; Faculty of Medicine and Health Sciences, Tel Aviv University, Tel Aviv, Israel; Division of Renal Medicine, Brigham and Women’s Hospital, Boston, MA, USA; Harvard Stem Cell Institute, Harvard Medical School, Boston, MA, USA

## Abstract

The kidney maintains homeostasis through an array of parallel nephrons, which all originate in development as isolated epithelial structures that later fuse through their distal poles to a system of collecting ducts (CD). This connection is required to generate functional nephrons by providing a pathway for excretion of metabolic waste and byproducts. Currently, methods for differentiating human pluripotent stem cells into kidney organoids generate nephrons that lack CDs and instead terminate as blind-ended tubules. Here we describe a developmentally inspired system that addresses this deficiency through assembly of induced nephrogenic mesenchyme with ureteric bud (UB) tissues, the embryonic building blocks of the kidney’s collecting system. The UB progenitors grow and develop into a network of CDs within the organoid, and importantly, they functionally integrate with the nephrons through recapitulating fusion between the distal tubule and CD to create a continuous epithelial lumen. We further showed that proximal-distal nephron specification, fusion frequency, and maturation of the CD can be augmented through temporal manipulation of developmental signaling pathways. This work provides a platform for interrogating the principles and mechanisms underlying nephron-UB fusion and a framework for engineering unobstructed nephrons with patterned collecting systems, an important step toward the de novo generation of functional kidney tissue.

## Introduction

Generating tissues from human pluripotent stem cells (hPSCs) capable of replicating the diverse and complex physiology of the mammalian kidney depends on recapitulating key structural features that are established during organogenesis. Each nephron (of up to 1 million in the human kidney) comprises a stereotyped series of specialized tubules that eventually reaches the collecting duct (CD) system, which transports formative urine through the corticomedullary axis to the ureter for elimination. Although nephrons and their collecting system are seamlessly connected to work in concert with one another in the adult kidney, they originate in the embryo from two distinct epithelial populations derived from separate progenitor pools: nephron progenitor cells (NPCs) and the ureteric bud (UB)^1–3^, respectively. At an early stage of nephrogenesis, the separate epithelia are permanently connected via a naturally-occurring anastomotic junction that establishes a patent luminal conduit and reinforces the proximal-distal polarity of the future nephron^4,5^, and this fusion process is thus among the most critical developmental determinants of kidney function.

Current protocols for differentiating kidney organoids from hPSCs rely on directed specification of nephrogenic mesenchyme (NM), which includes NPCs that can be induced to epithelialize and form nephron-like structures comprising podocytes, proximal tubules, and distal-like tubules^6,7^. Despite advances in these methodologies^8–10^, a persistent limitation has been that organoids lack CDs or any form of a collecting system that would provide a potential structural mechanism for the distal drainage of fluid from their nephrons. This is a fundamental barrier to advancing the maturation of organoids since distal tubular obstruction excludes the possibility of renal function. Additionally, the collecting system serves as an important centralized structure providing organization to the otherwise chaotic-appearing renal cortex. Though nephrons are densely and randomly arranged with respect to one another, they all connect to CDs through their distal ends to generate a uniform directionality and tubular axis that maximizes the function of the organ. The nephrons within organoids are similarly disordered but do not contain CDs to choreograph their orientation or collective action. The absence of a collecting system in kidney organoids is therefore a critical limitation and resolving this issue will be an essential step toward *de novo* production of physiologically competent human kidney tissue.

The UB generates the collecting system of the kidney, and it arises from an anatomically distinct region of the embryo compared to the NM in the metanephric mesenchyme^2,11^. UB progenitors are thus not induced in standard kidney organoid protocols^12,13^, so separate strategies for differentiation of this population have emerged in recent years^14–16^. Proof-of-concept studies using mouse embryonic stem cells showed that combinations of induced UB and NM could reproduce key developmental interactions from the nephrogenic niche and generate remarkably well-patterned collecting systems *in vitro*^14,17^. However, attempts to combine hPSC derivatives have not yielded comparable results. Mixing dissociated single cells from NPC and UB-like lineages led to rare chimeric epithelial structures^12,13,18^, but neither organized nephron-UB fusion nor the formation of CD-like tubules that could potentially drain organoid nephrons have been achieved. We recently reported the development of hPSC-derived UB progenitor cells that exhibited the potential to undergo branching morphogenesis and maturation to CD epithelia when cultured in isolation^16^. Here we describe a system for assembly of these UB progenitors with NM to incorporate CD-like drainage tubules into kidney organoids, and this system robustly recapitulates epithelial fusion between distal nephron and CD that parallels the process observed during *in vivo* development.

## Results

### Assembling hPSC-derived UB and NM progenitors into kidney organoids

To introduce a collecting system in kidney organoids, we sought to reconstruct the human nephrogenic niche through combination of UB^16,19^ and NM^7^ progenitors induced from parallel hPSC directed differentiation protocols (as summarized in Fig. 1A and Supp. Fig. 1A and 1C). To distinguish the distinct lineages in co-culture, hPSCs constitutively expressing GFP (from the permissive *AAVS1* locus) were used to generate UBs that were combined with GFP^-^ NM cells. To maximize potential developmental interactions, we dissociated NM at day 8, which was enriched for SIX2-expressing NPCs (Supp. Fig. 1B), mixed them with intact day 6 UB spheroids that largely comprised RET^+^ tip-like progenitors^16^ (Supp. Fig. 1D), and then cultured the aggregated tissue mixtures on transwell membranes as shown in Fig. 1A. Given the temporal dyssynchrony, we reset the differentiation numbering to be day 0 on the day of mixing the progenitors, and all subsequent reference to staging is relative to this timeframe. Though individual NM and UB organoids require different signaling conditions in isolation, we empirically optimized a simplified protocol to promote tissue interactions while still supporting the differentiation of these two lineages. Transient exposure to ROCK inhibitor (Y-27632, 10 uM) for the first 5 hours post-mixing enhanced the aggregation efficiency and augmented the differentiation of the NM lineage (Fig. 1B and Supp. Fig. 2A). In addition, BMP inhibition (LDN193189, 200 nM) from days 0-2 further improved the differentiation, as indicated by renal vesicle formation at day 4 (Supp. Fig. 2B). Beyond day 2, the co-culture differentiation occurred under permissive conditions in basal medium supplemented with 10% serum. Importantly, this method entirely avoided the use of GSK3β inhibition to mimic WNT-induced NM epithelialization, which was omitted given its likely effect in biasing subsequent nephron differentiation^20,21^.

**Figure 1.**
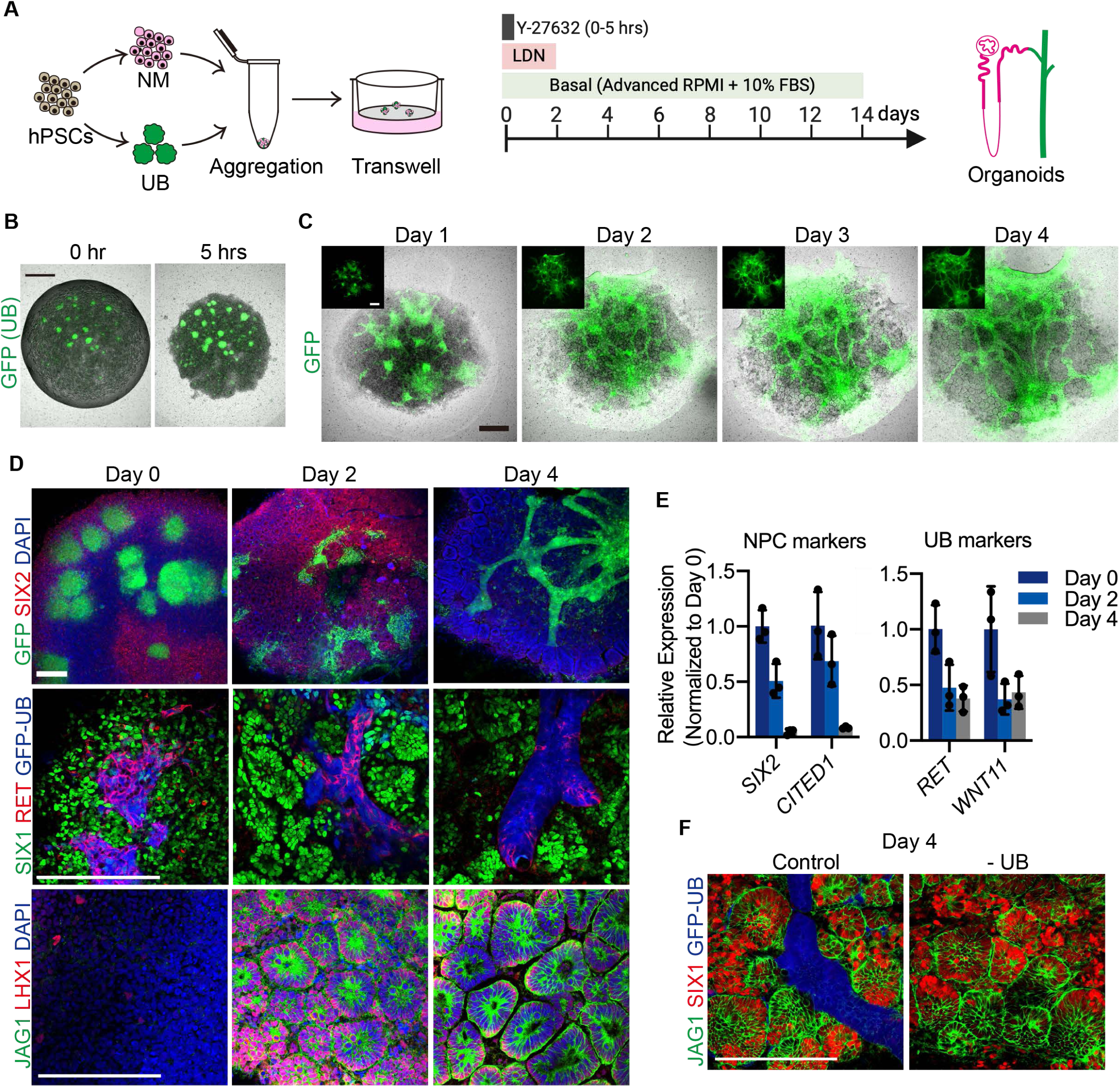
Assembly and dynamics of UB and NM progenitor cells in integrated kidney organoids. **A.** Schematized protocol for integration of induced UB and NM into kidney organoids on transwell membranes. **B.** Within 5 hours, the dissociated NM condensed into a disk-shaped structure and surrounded the embedded GFP^+^ UB spheroids. **C.** In several days, the NM was induced into epithelialized renal vesicle-like structures, while the GFP^+^ UBs grew as elongating and sometimes bifurcating tubular epithelia that penetrated throughout the organoid. Insets show the isolated GFP channel. **D.** Whole-mount IF staining revealed progressive loss of NM progenitor markers SIX1/2 from days 0-4 with concomitant nearly uniform induction of renal vesicle markers LHX1 and JAG1 accompanying adoption of epithelial morphology. The GFP^+^ UBs exhibited similar reduction of the tip progenitor gene RET, although patchy expression was still observed on day 4. **E.** qPCR analyses confirmed loss of undifferentiated NPC markers *SIX2* and *CITED1* from day 0 to day 4 and reduction of UB tip progenitor markers *RET* and *WNT11*. *n*=3 organoid replicates per timepoint. **F.** NM progenitors displayed similar differentiation patterns when cultured in the absence of UB spheroids. Scale bars, 500 μm (B-C) and 200 μm (D, F).

Following aggregation, the GFP^+^ UB spheroids embedded within NM progenitors expressing undifferentiated markers such as SIX2 and SIX1 (Fig. 1D), reminiscent of the cortical nephrogenic niche in developing kidneys. However, within 48 hours we observed widespread nephron induction characterized by formation of epithelialized vesicles expressing LHX1 and JAG1 (Fig. 1C-D), and by day 4 undifferentiated progenitors were largely undetectable (Fig. 1D-E). Meanwhile, despite the loss of associated NPCs, the recombined UB spheroids grew extensively during this period to form a network of tubules interwoven amongst the nascent nephrons throughout the organoids (Fig. 1C). The early stages (days 0-2) of their morphogenesis involved the rapid formation of numerous buds from each individual spheroid that subsequently underwent further sprouting or branching, with later growth (beyond day 3) predominantly consisting of tubular elongation. We have observed similar patterns using UB organoids derived from multiple different hPSC lines, including *GATA3*- mScarlet reporter H9 cells (Supp. Fig. 2C).

The rapid epithelialization and exhaustion of progenitors was consistent with previous reports describing a single wave of differentiation within kidney organoids rather than the normal iterative process that occurs *in vivo*^22^. The same pattern was observed in our system in the absence of UBs (Fig. 1F and Supp. Fig. 2D), indicating that they were not sufficient to alter the differentiation behavior of NPCs. In contrast, molecular analyses revealed that tip progenitor markers, such as *RET* and *WNT11*, indeed persisted within the UB epithelia until at least day 4 (Fig. 1D-E), excluding their absence as the cause for the failure to recreate the self-renewing niche. We hypothesized insufficient signaling capabilities between the two compartments as an alternative etiology, but the addition of exogenous FGF and GDNF, aimed to promote maintenance of NPC and UB progenitor states^23,24^, respectively, did not overtly impact development in the organoids (Supp. Fig. 2E). Thus, the inclusion of UB spheroids into kidney organoids did not significantly impact the NM lineage at the progenitor stage, but it did successfully generate an integrated network of tubules that could potentially serve as a CD-like drainage system.

### Distal nephron segments fuse to UB-derived CDs through a developmentally conserved process

The recombinant organoids were grown on the transwell filters under permissive conditions, and by day 14 they exhibited several remarkable features that have not previously been reported in hPSC-derived kidney tissues. First, while the GFP^-^ NM developed into a dense collection of differentiated nephron epithelia as previously described^6,7^, the UB progenitors had grown into elongated CD-like tubules that were largely interconnected and spanned throughout the organoids (Fig. 2A). Also notable in the images was the expansion of a GFP^+^ stromal compartment that is otherwise not extensively characterized as part of this study. The formation of expected nephron segments, including podocytes, proximal tubules, and distal tubules, was not markedly affected by the co-differentiation with UB, although it was associated with a slight reduction in podocyte markers (*NPHS1*) measured by qPCR (Fig. 2C). In contrast, the expression of distal/CD markers *GATA3*, *CALB1*, *AQP2*, and *SCNN1B* was significantly increased (Fig. 2C), consistent with the development of UB progenitors into CD tissues. *AQP2*, for example, was largely undetectable in NM organoids and elevated approximately 90-fold in the mixed lineage organoids. Most interesting, though, was the observation that numerous GFP^-^ nephron tubules were directly connected to the GFP^+^ ducts (Fig. 2B), and this appeared morphologically analogous to the fusion of the nephron connecting tubule to the CD. In NM-only organoids, there were rare putative connecting tubules expressing CALB1^25^, but they were short, blunted segments that blindly terminated as dead ends (Fig. 2D). In contrast, recombinant organoids contained extended CALB1^+^ distal nephron segments that connected with GFP^+^ CDs that also expressed CALB1 (Fig. 2D). Closer examination through wholemount staining and confocal imaging confirmed the presence of *bona fide* fusion between nephron segments and CDs. Evaluation of apical (TJP1 and PRKCZ, also known as ZO1 and aPKC, respectively) and basal (CDH1 and Laminin) polarity markers revealed an uninterrupted epithelial structure and continuous luminal membrane across the junctions of GFP^-^ and GFP^+^ tubules (Fig. 2E and Supp. Fig. 3A). Overall, the epithelial fusion was a common and highly reproducible event, occurring numerous times (typically dozens, as quantified below) in every organoid we have examined (>500).

**Figure 2.**
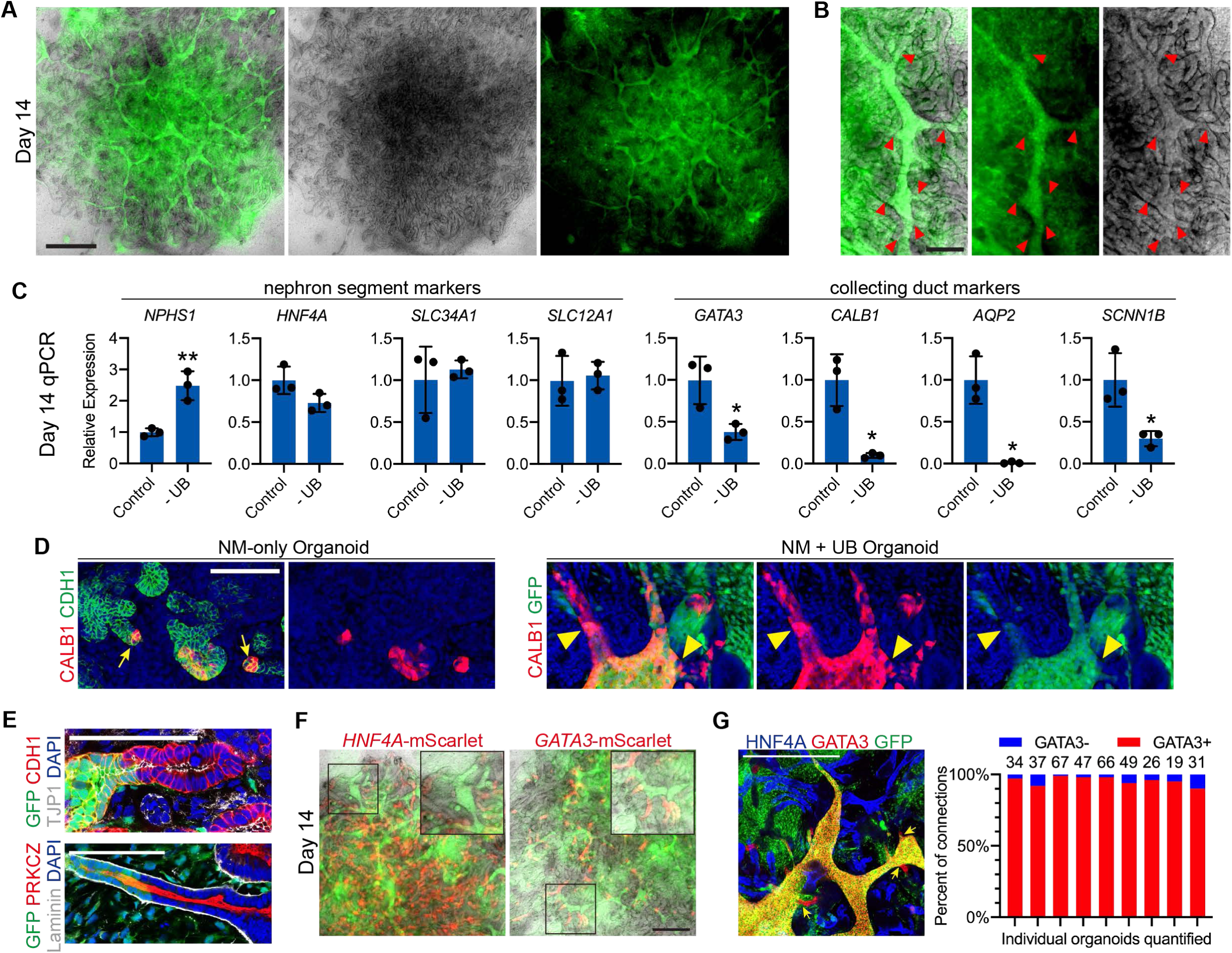
Formation of collecting ducts in organoids that fuse to distal nephron tubules. **A.** Following two weeks of culture, the GFP^+^ UBs generated extensive networks of CD tubules that were embedded amongst the nephron epithelia of the organoid. **B.** Numerous points of epithelial fusion (marked by red arrowheads) were observed between the GFP^+^ CDs and unlabelled nephron tubules. **C.** Gene expression analysis by qPCR at day 14 revealed lower levels of *NPHS1* (\*\**P*J=J0.0059) but otherwise comparable expression of nephron segment differentiation between mixed (Control) organoids and those without UB. Control organoids had significantly upregulated CD differentiation markers *GATA3* (\**P*J=J0.0236), *CALB1* (\**P*J=J0.036), *AQP2* (\**P*J=J0.0261), and *SCNN1B* (\**P*J=J0.0214). *n*=3 independent organoid replicates per group; column and error bars represent mean and standard deviation, respectively. **D.** Expression of CALB1 in NM-only organoids marked the presumptive connecting segments that terminated as blind-ended tubules (yellow arrows), while in integrated organoids these segments fused to GFP^+^ CDs also expressing CALB1 (yellow arrowheads indicate epithelial fusion points). **E.** Nephron-CD anastomoses at day 14 exhibited uninterrupted apicobasal polarity with apparent continuity of the apical lumen across the junction. **F.** Organoids at day 14 contained an abundance of *HNF4A*^+^ proximal tubules and a relative scarcity of *GATA3*^+^ distal segments, as shown in micrographs of live organoids harboring fluorescent reporter alleles. The UBs are shown in green (GFP). **G.** IF staining and image quantification revealed that 96% of epithelial connections (as shown by yellow arrows) involved a GATA3-expressing nephron tubule, and fusions with HNF4A^+^ proximal tubules were not observed. Scale bars, 1,000 μm (A), 200 μm (B), 100 μm (D-E), and 500 μm (F-G).

*In vivo,* developing nephrons fuse to the UB through their distal-most segments that express GATA3^26^ and are adjacent to the UB. Prior work showed that specification of the distal tubule was required for fusion^4^. We therefore asked whether this phenomenon was preserved in kidney organoids, which do not exhibit the same high degree of organization and stereotypic spatial relationships. Using NM harboring fluorescent reporter alleles for proximal tubule (*HNF4A*; Supp. Fig. 4A) and distal tubule (*GATA3*^16^) identity, we found that the nephron epithelia comprised mostly proximal tubules and only a comparatively very minor population of *GATA3*^+^ distal segments (Fig. 2F). The same result was confirmed via wholemount staining (Supp. Fig. 3B), although this method also included the expected expression of GATA3 in the UB epithelium. Yet despite their vastly disproportionate under-representation, fusion of *GATA3* (mScarlet)-expressing tubules to GFP^+^ UB ducts was readily observed in the organoids (Fig. 2F-G and 3A). In many cases, multiple *GATA3*^+^ tubules were connected to a larger GFP^+^ duct, suggesting that the UBs developed into central CD-like structures capable of draining numerous nephrons in a given area. Quantification of confocal images showed that each organoid contained a mean of 41.8 ± 16.8 (s.d.) epithelial connections between nephron and UB. Astoundingly, 96% of the fused nephron segments expressed GATA3 (Fig. 2G), and we have yet to observe connection between an HNF4A^+^ proximal tubule and the UB. These data therefore support the presence of robust mechanisms that constrain the fusion competence of developing renal epithelia, which is essential to establish a normal proximal-distal axis in the nephron, and they are conserved in the kidney organoids irrespective of the random orientation of nephrons in this system.

Having observed that fusion was specific to the distal tubule, we further characterized the temporal development of nephron segmentation and UB connection in the organoids. As described above (Fig. 1C), the NM progenitors robustly formed epithelialized structures resembling renal vesicles by day 4. At this stage, the vesicles exhibited evidence of early polarization with WT1 and POU3F3 marking the presumptive proximal and distal domains, respectively (Fig. 3C), although there was still significant overlap of expression. As early as day 4-5, *GATA3* expression was initiated as a small patch of cells in a portion of the vesicles (Fig. 3A-B), which between days 5-7 expanded into a short primitive connecting segment. As shown in daily images of live organoids in Fig. 3B, when this region formed adjacent to a UB the *GATA3*^+^ progenitors appeared to interact and invade into the GFP^+^ epithelium. This process involved the extension of the *GATA3* domain toward the CD, along with its apical membrane, which then quickly established a continuous apical luminal surface with that of the CD (Fig. 3D-E). Correspondingly, the basement membrane that encapsulated the renal vesicles appeared to have been broken down at the sites of fusion (Fig. 3E). Overall, the process was surprisingly synchronous among the many nephrons in the organoids as it primarily occurred within a 2-3 day window. By day 7, most of the fusion events were completed and the organoids largely comprised well-segmented nephrons, a subset of which were fused to UB-derived CDs (Fig. 3F and Supp. Fig. 3C).

**Figure 3.**
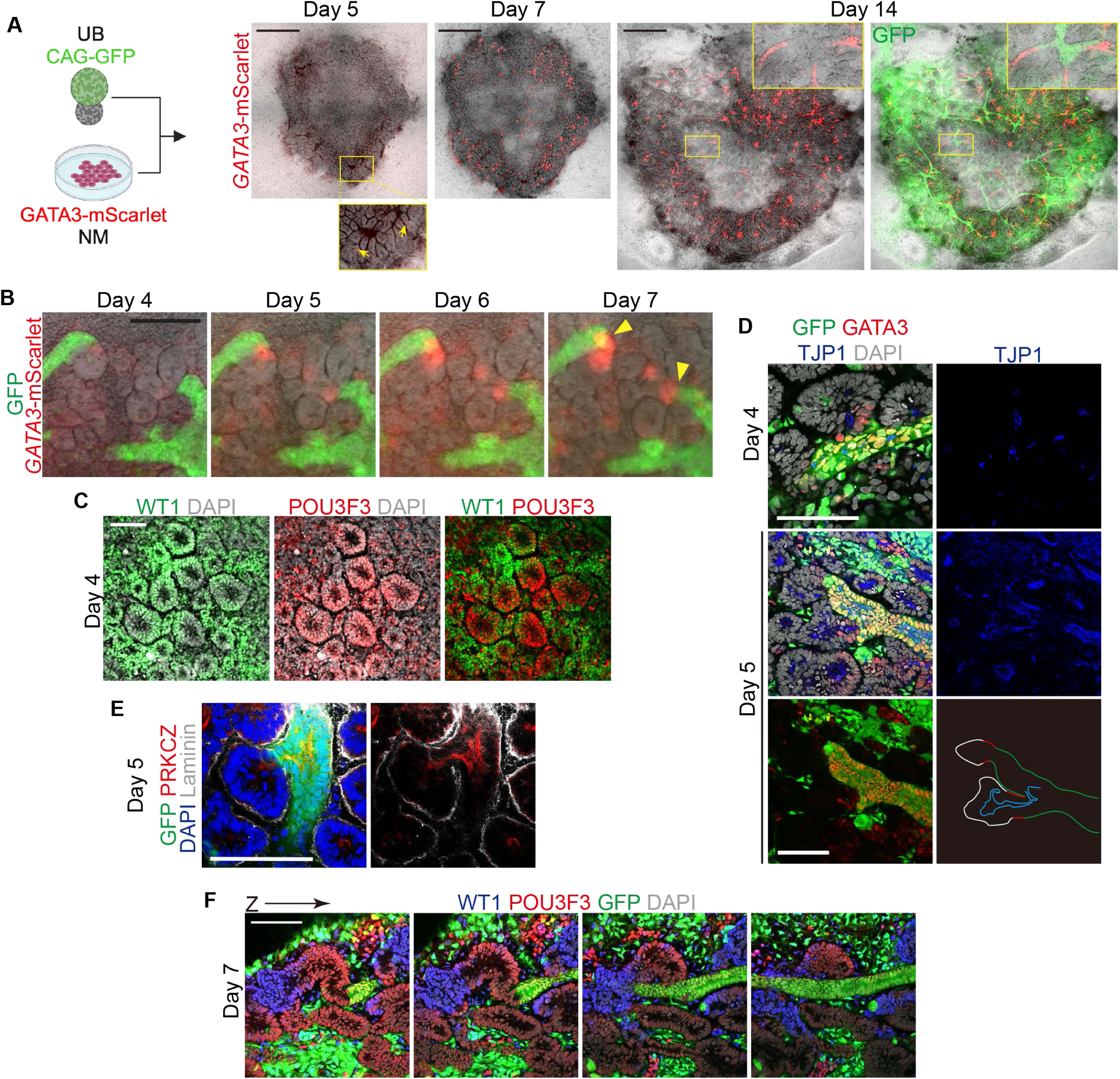
Fusion to the UB follows nephron polarization and segmentation. **A.** *GATA3*-mScarlet expression in the nephron lineage was first weakly detected as early as day 5 in small domains of renal vesicles (yellow arrows), and by day 7 it was strongly expressed in the presumptive distal segments of nascent nephrons. By day 14, the distal tubules were frequently fused to the GFP^+^ CDs. **B.** Daily imaging revealed the process by which the early *GATA3*^+^ segment interacts with and invades into nearby GFP^+^ UB epithelia. Yellow arrowheads indicate points of nephron fusion. **C.** At day 4, the renal vesicles exhibited polarization with coarse segregation of the proximal (WT1) and distal (POU3F3) domains. **D.** GATA3 expression in these early polarized vesicles was often associated with extension of the epithelium and its apical membrane (TJP1) toward the UB, and complete apical connections were observed by day 5. **E.** Loss of extracellular matrix (Laminin) was observed at the site of fusion while the apical membrane (PRKCZ) extended across the junction between the renal vesicle and UB. **F.** Optical sections through an organoid at day 7 revealed segmented and sometimes organized nephrons progressing from WT1^+^ presumptive glomerular structures through the distal epithelial fusion with the UB. Scale bars, 1,000 μm (A), 200 μm (B), and 100 μm (C-F).

### Notch inhibition augments tubule fusion via nephron distalization

One intriguing observation from the above experiments was that only a subset of the renal vesicles ever formed a *GATA3* domain, and those without one seemed incapable of interacting and fusing with the UB epithelium (Fig. 3B). Combined with the apparent exclusivity of the anastomoses to this relatively rare population (Fig. 2G), this led us to hypothesize that the overall frequency of nephron-CD fusion could be augmented by improving the efficiency of specification of these distal nephron segments. Given its role in promoting proximal nephron formation^27–29^, we tested whether inhibition of NOTCH signaling (using the gamma secretase inhibitor DAPT, 10 uM) could be strategically used to manipulate the proximal:distal differentiation ratio. The organoids were exposed to DAPT for varying lengths of time between days 2-6 (Fig. 4A), the period in which molecular patterning and segmentation of the nascent nephrons was occurring. Remarkably, prolonged NOTCH inhibition between days 2-6 induced nearly complete distalization of the nephrons at day 8, with >99% reduction in HNF4A^+^ proximal tubular area and a corresponding 39-fold increase in GATA3^+^ tubules (Fig. 4B). By day 14, this led to markedly abnormal-appearing organoids that contained mostly amorphous GATA3^+^ epithelial structures, which resulted from fusion both between the vesicles and with the GFP^+^ UBs, and a paucity of proximal tubules and podocytes (Fig. 4C and Supp. Fig. 5A).

**Figure 4.**
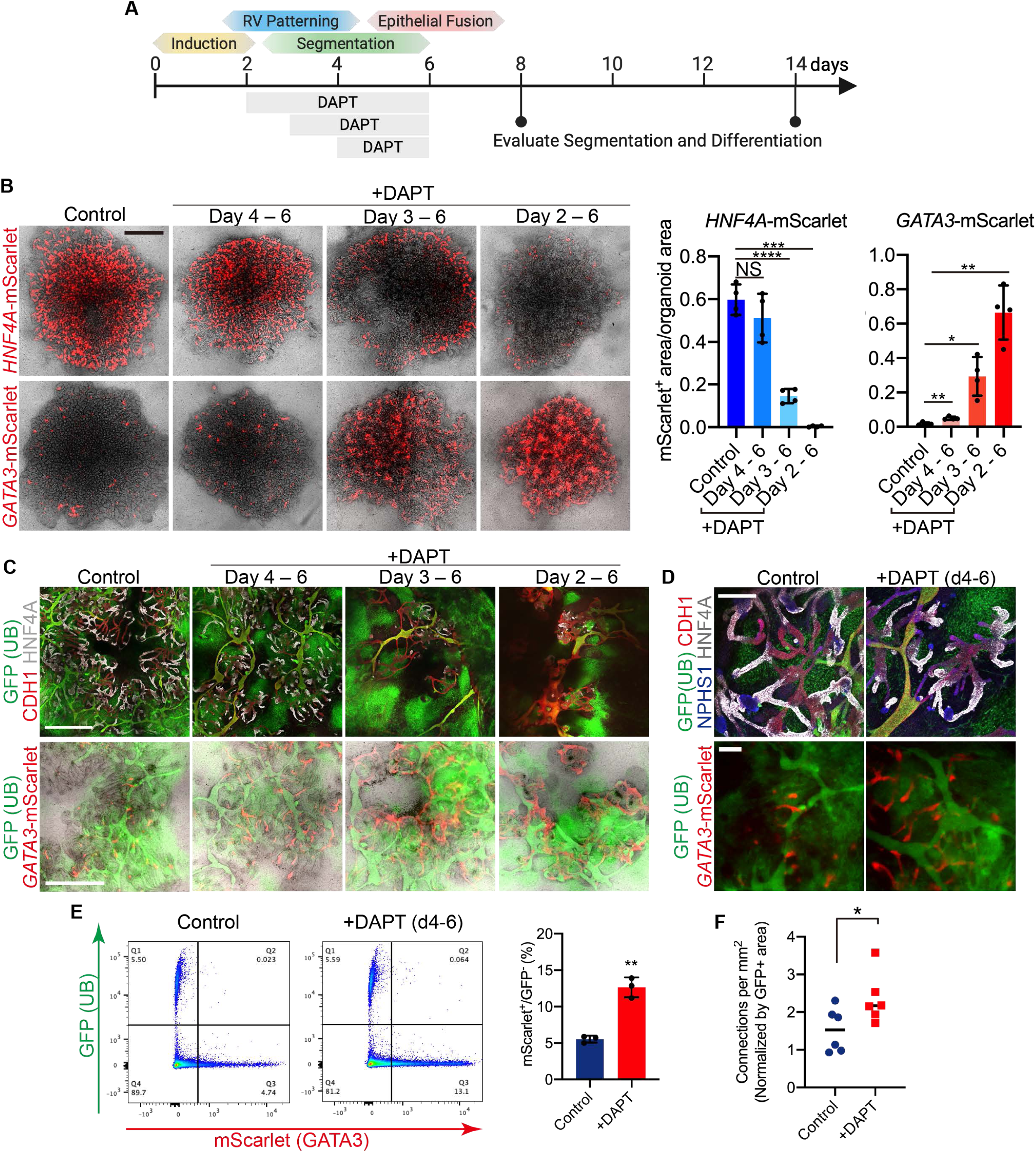
NOTCH inhibition augments distal nephron specification and fusion competence. **A.** Summary of early nephrogenesis events in organoids (day 0 = time of mixing NM and UB) and strategy for testing temporal inhibition of NOTCH signaling. **B.** Proximal (*HNF4A*) and distal (*GATA3*) nephron specification visualized and quantified in live organoids at day 8 via mScarlet fluorescent reporter activity. Prolonging exposure to the NOTCH inhibitor DAPT during the segmentation stage led to progressive reduction of proximal and increase in distal nephron formation. *n*=4 independent biological replicates per condition; \*\*\*\**P*J<0.0001, \*\*\**P*J=J0.0005, \*\**P*J=0.0024 (control vs. day 4-6), \*\**P*J=J0.0037 (control vs. day 2-6) and \**P*J=J0.016. **C.** At day 14, nephron epithelia displayed disorganized morphology in organoids treated with 3-4 days of DAPT with widespread *GATA3* expression and loss of HNF4A^+^ proximal tubules, whereas shorter treatment (days 4-6) led to increased abundance of the short *GATA3*^+^ distal segments compared to controls but with preserved overall morphology and maintenance of proximal tubular development. **D.** The organoids treated with DAPT from days 4-6 retained a comparable number of HNF4A^+^ proximal tubules and NPHS1^+^ podocytes, and they contained more *GATA3*^+^ distal segments that were fused to the UB-derived ducts. **E.** DAPT treatment induced 2.5-fold expansion in *GATA3*^+^ distal nephron cells by flow cytometry (*n*=3 independent biological replicates per condition; \*\**P*J=J0.001) and (**F**) significantly increased frequency of nephron-UB fusion events (*n*=6 independent organoids per condition; \**P*J=J0.045). Scale bars, 1,000 μm (B, C), and 200 μm (D). Column and error bars represent mean and standard deviation, respectively.

Supporting the dynamic nature of the developmental events during organoid formation, the DAPT-induced distalization was exquisitely sensitive to the timing and duration of exposure. Shortening the treatment to 3 days (3-6) produced a similar yet less severe phenotype with a 76% decrease and 17-fold increase in *HNF4A* and *GATA3* tubules at day 8, respectively. Further reduction to just a 2-day pulse (4-6) led to only a modest (and non-significant) 14% reduction in *HNF4A* (Fig. 4B), indicating that proximal specification was mostly established by day 4 and was irreversible. Conversely, DAPT from days 4-6 did cause a beneficial effect on the formation of the *GATA3*^+^ connecting tubule, with a 2.9-fold increase in *GATA3*^+^ area at day 8 (Fig. 4B) and a 2.5-fold increase in *GATA3*^+^ nephron cells at day 14 (Fig. 4C-E). We further confirmed the preservation of podocyte, proximal tubule, and thick ascending limb differentiation in this condition (Fig. 4C-D and Supp. Fig. 5A-B), which would be important to ultimately generating fully segmented nephrons.

Accompanying the increased specification of *GATA3*^+^ nephron segments, there was increased frequency of fusion events with the UB (Fig. 4C-D and Supp. Fig. 5A). Quantification showed a significant increase in the total number of anastomoses per organoid, which was normalized to the amount of UB tubules, following treatment with DAPT from days 4-6 (Fig. 4F). We confirmed that these epithelial connections exhibited the same properties as we previously characterized, including continuity of the apical membrane across the junction and expression of CALB1 in the NM-derived connecting segment (Supp. Fig 5C-D). Collectively, these data further support the conclusion that distal nephron specification has a deterministic role in promoting the ability of tubules to fuse to the CD, and NOTCH perturbation may be used to alter the formation of fusion-competent distal tubules in organoids and consequently, the frequency of anastomoses.

### Coordinated and parallel development of UB and nephron lineages in co-culture

Single cell RNA-seq analyses were performed to further define the lineage compositions and developmental trajectories within the recombinant kidney organoids at days 3, 7, and 15. Following demultiplexing, removal of multiplets, and filtering of low-quality cells, the combined dataset of 11,330 cells revealed a high level of complexity in the organoids with 16 unique clusters comprising nephron, ureteric, stromal, and endothelial populations, which we identified via expression of canonical marker genes (Supp. Fig. 6A-E). As anticipated, the UB lineage label GFP was expressed nearly uniformly (>93%) in cells of the ureteric clusters and largely excluded (∼3%) from the nephron lineage (Supp. Fig. 6F). GFP^+^ cells also contributed to 58% of the stromal population and 29% of the small endothelial cluster (Supp. Fig. 6F), representing likely expansion and differentiation of the interstitial progenitor cells present in the UB spheroids at day 6 (Supp. Fig. 1D). To focus subsequent analyses, the stromal and endothelial clusters (Supp. Fig. 6G) were removed to isolate the 6,834 cells representing UB and NPC development.

Supervised annotation of the re-clustered dataset using established anchor genes showed 8 clusters representing stages of nephron lineage differentiation and a seemingly distinct population comprising 2 clusters with UB-like transcriptional profiles (Fig. 5A and Supp. Fig. 7A-B), which were labelled as ‘Early’ and ‘Late.’ We further evaluated these clusters through unsupervised analyses using DevKidCC^30^, which corroborated the distinct ‘Nephron’ and ‘Ureteric Epithelial’ lineages within the organoids in the ‘Tier 1’ analysis, as well as subsequent characterization of nephron segmentation (Fig. 5B). A small subset of the day 3 cells was predicted as ‘NPC’-like by DevKidCC, but these cells did not exhibit *SIX2* expression (Supp. Fig. 7A), consistent with the rapid exhaustion of undifferentiated NPCs we observed by day 2 (Fig. 1D-E). In contrast to the iterative nature of *in vivo* kidney development, the scRNA-seq analyses further demonstrated that differentiation within organoids was remarkably well synchronized in a single wave of nephron formation. Each of the clusters in the nephron lineage was composed of a dominant timepoint representing >80% of the cells (Fig. 5C), except for the LOH (73% from day 15, 27% from day 7). Cells at day 3 were mostly in an early renal vesicle-like state (PTA/RV) that later segregated by day 7 into proximal, distal, and podocyte-like progenitor populations analogous to those found in the SSB, consistent with our earlier analyses (Fig. 3). Differentiation into more mature epithelial segments and cell types was observed at day 15.

**Figure 5.**
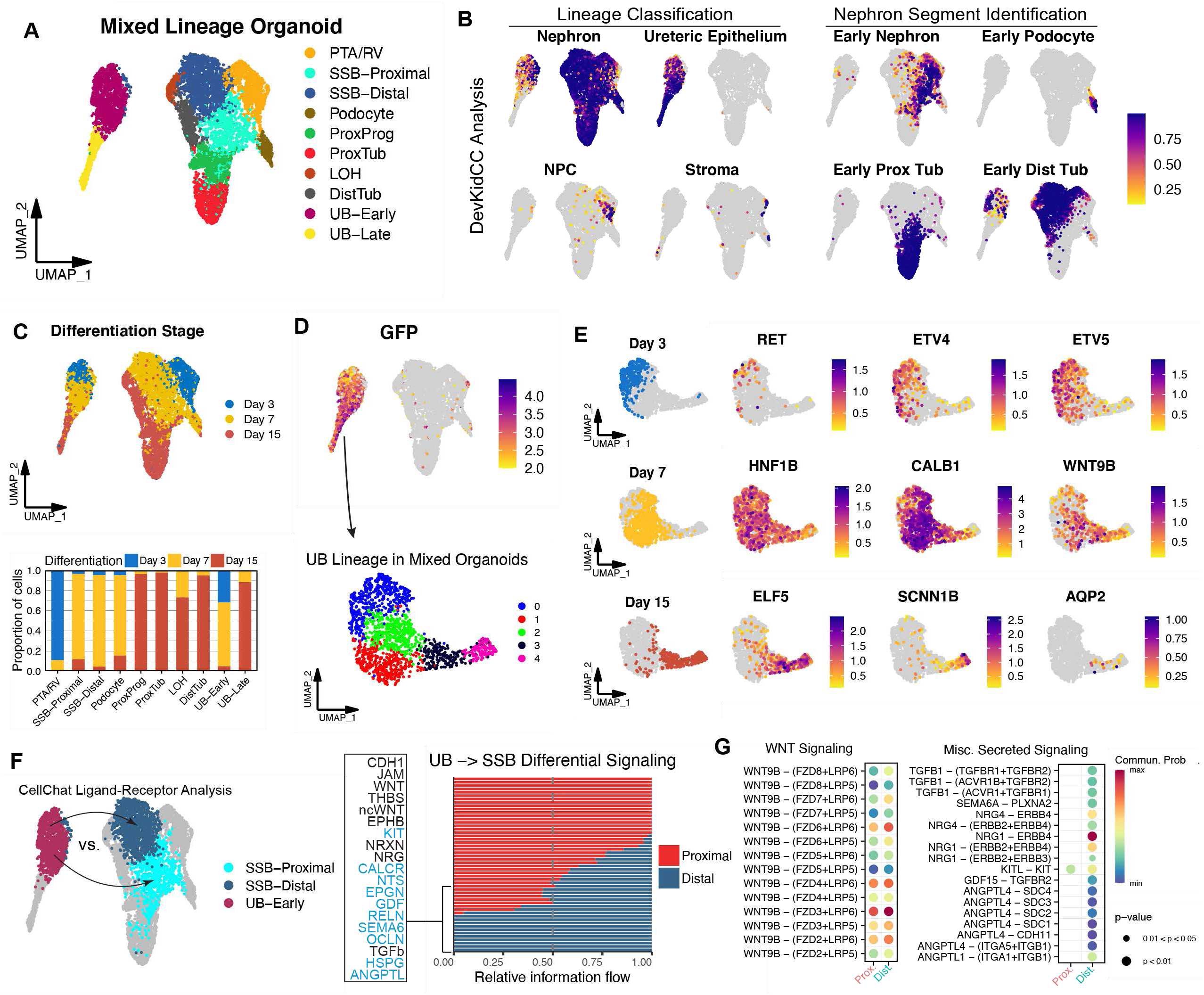
UB and NM progenitors are lineage-restricted and differentiate in parallel. **A.** UMAP embedding of recombinant kidney organoids with supervised cluster annotation. **B.** DevKidCC assignment scoring for lineage classification (Tier 1) and nephron segmentation (Tiers 2/3). **C.** Cells in the dataset largely segregated by their stage of differentiation, days 3, 7, and 15. **D.** Expression of the lineage label GFP was specific to the two UB clusters, confirming the lineage fidelity of the early progenitors. These cells were extracted and re-analyzed to reveal 5 clusters representing the differentiation trajectory of the UB lineage. **E.** The UB cells at day 3 were enriched for tip/progenitor markers, while cells on day 15 exhibited a more mature CD signature. Day 7 cell exhibited an intermediate phenotype including expression of the stalk progenitor marker *WNT9B*. **F.** A CellChat analysis was performed to compare the putative signaling interactions that originate in the UB and are enriched in distal over proximal receiver probability. Pathways in blue show statistically significant (*P* < 0.05) increases in signaling probability. **G.** Bubble plot showing predicted increased receptivity of the early distal tubule for UB-derived *WNT9B* signaling, as well as other secreted factors including TGFB and NRG family members.

Expression of GFP further confirmed that the UB and NM progenitors exhibited lineage-restricted potential at the time of aggregation, as it was highly expressed in the 2 UB clusters of the dataset and mostly restricted from the nephron lineage clusters (Fig. 5D). There was a small group of GFP^+^ cells in the PTA/RV cluster, but they had negligible scores for either ‘NPC’ or ‘Nephron’ lineages and were instead predicted as ‘Stroma’ by DevKidCC (Fig. 5B), suggesting they were likely UB spheroid-derived stromal progenitors. Closer examination and re-clustering the GFP^+^ UB clusters showed that they similarly progressed through developmental stages in a linear fashion (Fig. 5D-E). Cells at day 3 were enriched for tip-like progenitor genes including *RET*, *ETV4*, and *ETV5*, while at day 7 there was more expression of *CALB1* and *WNT9B* that are associated with the UB stalk fate. By day 15, most of the UB-derived cells exhibited signatures consistent with CD principal cells including the transcription factor *ELF5*, sodium channel subunit *SCNN1B*, and water channel *AQP2*. Further investigation of the maturation of this lineage is discussed below.

We sought to determine whether the molecular profiling could identify potential candidates that might be mechanistically involved in the epithelial fusion process between the UB and distal nephron. CellChat analysis was performed to specifically interrogate predicted signaling interactions between the UB-Early, SSB-Distal, and SSB-Proximal clusters (Fig. 5F), which largely comprised cells from day 7 of differentiation when fusion was ongoing (Fig. 3B-F). Enriched ligand-receptor pairings from the UB to distal tubule, which might underlie the unique fusion potential of this segment, included both canonical (via WNT9B) and non-canonical WNT signaling, which both govern multiple aspects of kidney development^31–33^. In addition to these known candidates, many unexplored pathways emerged from this analysis, including secreted signals such as TGFβ, Semaphorins, and Neuregulins, among others (Fig. 5F-G). Ephrin signaling and other direct contact-mediated cell interactions, as well as potential communications through the ECM, were also enriched in the distal segment (Supp. Fig. 7C). This new integrated organoid system will serve as a robust platform for functionally testing these potential novel molecular regulators of fusion in future studies.

### Nephron-UB fusion occurs *in vivo* following organoid transplantation

Transplantation beneath the kidney capsule of immune-deficient mice provides a permissive environment for the vascularization, growth, and maturation of kidney organoids^34,35^. To determine whether the *in vivo* environment would also support the formation and maintenance of nephron-UB connections, we transplanted integrated organoids at day 3 prior to their *in vitro* fusion (Fig. 6A). By 2 weeks post-transplantation, the engrafted organoids had developed into complex tissues comprising both renal parenchyma and an expanded stromal compartment (Fig. 6B-C). Of note, the interstitial tissue included both GFP^-^ and GFP^+^ cells, and although the latter derived from UB organoids, they appeared to support the relatively dense and robust growth of nephron tubules (Fig. 6C). As anticipated, transplantation promoted the maturation of glomeruli at the proximal end of the nephron. Histological and immunofluorescent analyses revealed organized glomerular structures including arrayed podocytes (NPHS1), interstitial or mesangial cells (PDGFRB, GATA3), and endothelial cells (PECAM1) that formed capillaries containing erythrocytes (Fig. 6D). Distal to the glomeruli, many of the nephrons in the engrafted tissue exhibited appropriate segmentation (Fig. 6E) with sequential development of proximal tubules (HNF4A), loop of Henle segments (SLC12A1), and distal connecting tubules (GATA3).

**Figure 6.**
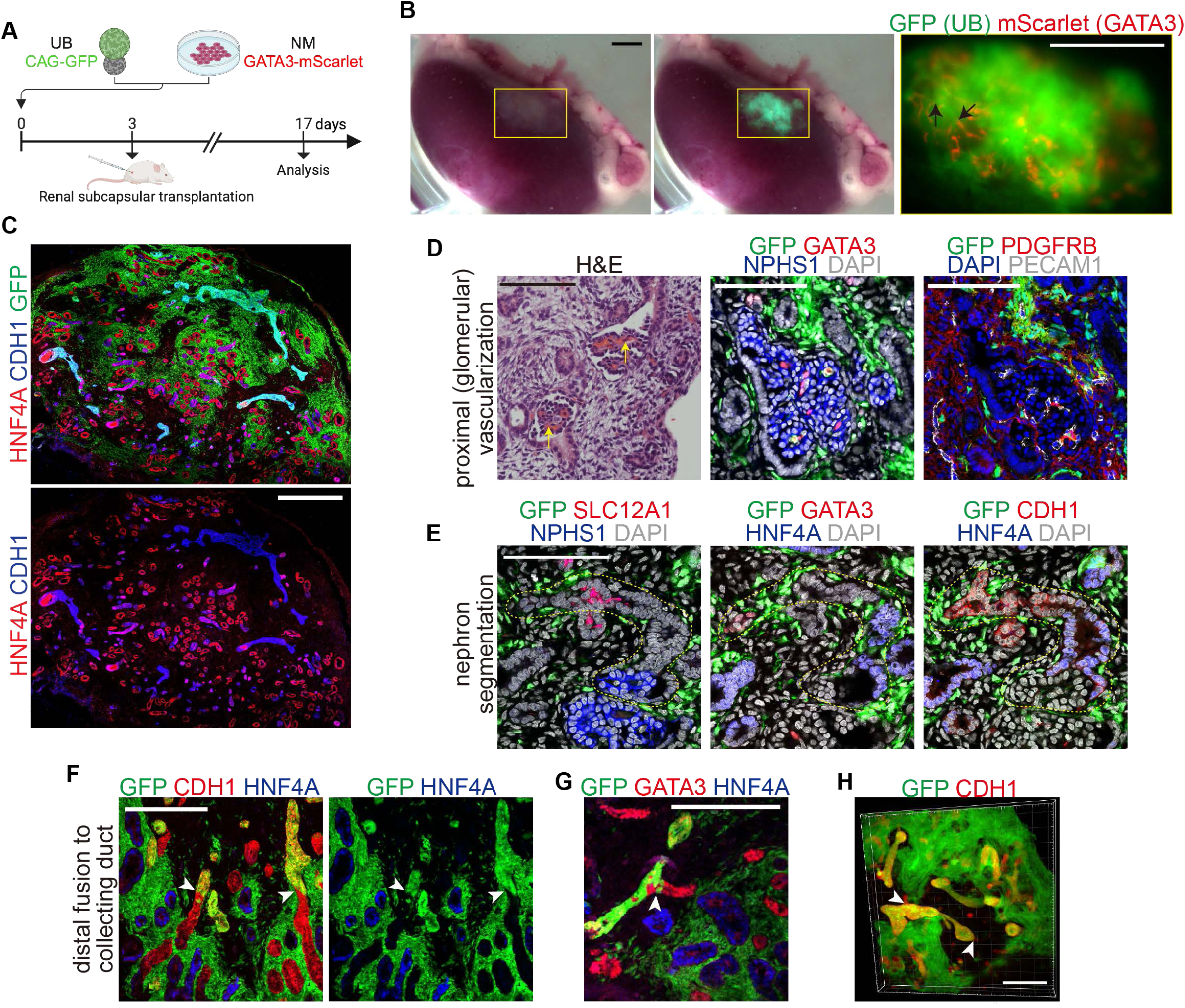
Fusion of UB and distal nephron following *in vivo* transplantation. **A.** Overview of organoid transplantation experiments. **B.** Gross appearance and stereomicrographs of organoid tissue on the kidney surface following two weeks of *in vivo* growth. Short *GATA3*^+^ nephron tubules were seen throughout the graft, including many that were connected to larger GFP^+^ duct-like structures (black arrows). The general haziness of the GFP signal indicated robust growth of UB organoid-derived stromal cells. **C.** The engrafted tissue comprised numerous NM-derived proximal (HNF4A) and distal (CDH1) tubules and large UB organoid derived duct structures and interstitial cells. **D.** *In vivo* growth enabled vascularization and maturation of organoid glomerular structures that comprised an organized arrangement of podocytes (NPHS1), endothelial cells (PECAM1), and mesangial cells (PDGFRB, GATA3). Apparent perfusion of the glomerular tufts was indicated by the presence of red blood cells (yellow arrows). **E.** Segmented arrangement of nephrons in the graft was confirmed through serial sections highlighting the sequential progression of podocytes (NPHS1), proximal tubule (HNF4A), thick ascending limb (SLC12A1), and connecting tubule (GATA3). **F-H.** Fusion of nephron tubules to GFP^+^ UB-derived ducts was observed in the engrafted organoids (white arrowheads) in both sections (F-G) and wholemount staining (H), and it was restricted to the CDH1/GATA3^+^ distal segments. Scale bars, 1,500 μm (B), 500 μm (C), 100 μm (D-E), 200 μm (F-G), and 300 μm (H).

Having established the expected proximal glomerular maturation and integration of nephrons in transplanted organoids, we further evaluated their distal ends. Using the lineage reporter system (Fig. 6A), numerous *GATA3*^+^ distal nephron segments were observed in the gross explant tissue, and we also observed larger GFP^+^ CDs in the transplanted organoids (Fig. 6B), although their visualization was partially obscured by the UB organoid-derived stromal cells. Even under low-power fluorescence microscopy, it was apparent that many of the *GATA3*^+^ tubules were directly terminating and connecting into the GFP^+^ CDs. Closer analysis with confocal imaging and wholemount staining confirmed this result (Fig. 6F-H). Numerous nephron-derived CDH1^+^ and GATA3^+^ tubules were fused to the UB-derived CDs. Like the observations *in vitro*, proximal nephron segments did not participate in these anastomoses. Thus, the developmentally conserved process mediating fusion between the nephron and UB could occur both *in vitro* and *in vivo*, which might have important implications for expanding future possibilities aiming to functionally incorporate hPSC-derived nephrons into host organisms.

### Improving the maturation of CD epithelia

Having established methods for the incorporation and structural integration of UB-derived CDs into kidney organoids, we further explored their maturation into functional cell types. Our scRNA-seq analyses showed that despite the progressive loss of progenitor genes and induction of those associated with a more differentiated CD state, only very few UB-derived cells at day 15 exhibited appreciable *AQP2* expression (Fig. 5E and Supp. Fig. 8A-C), which is both a canonical marker and a functionally critical channel in principal cells. This was a surprising finding given our previous report demonstrated robust spontaneous differentiation of the same UB progenitor cells into AQP2^+^ principal cells following a period of growth in 3D culture in isolation (Supp. Fig. 8D^16^). To exclude an inhibitory effect of the transwell culture format, we plated UB organoids alone on the membranes and grew them in otherwise the same conditions as described previously (UB Media). Using a new fluorescent reporter for *AQP2* (Supp. Fig. 4B), we showed that these tissues exhibited robust *AQP2* expression (Fig. 7A), whereas the same organoids grown in the ‘Mix Media’ (Fig. 1A) did not differentiate. Similarly, UBs grown in co-culture with NM activated the reporter when cultured in UB Medium but not in Mix Media (Fig. 7A), indicating that the culture medium was an important determinant of the differentiation potential of the UBs.

**Figure 7.**
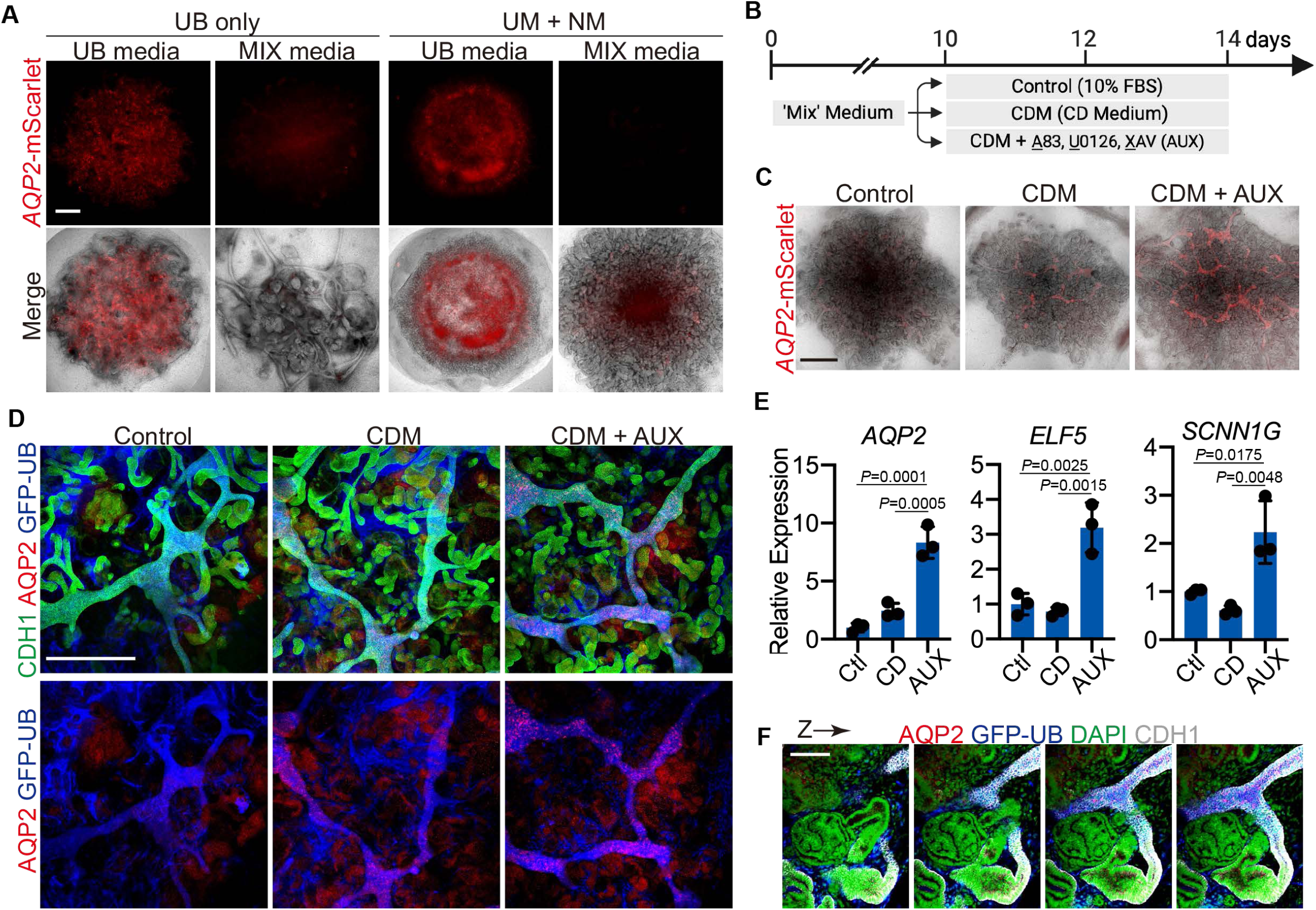
Induction of CD maturation in recombinant organoids. **A.** The expression of *AQP2* was induced in UB epithelia grown either in isolation or in recombinant organoids with NM when cultured in previously identified conditions to grow UB organoids (UB Medium), but not when grown in the minimal ‘Mix’ media. **B.** The combined organoids were transitioned at day 10 from ‘Mix’ medium to induce maturation. **C-E.** CD Medium (CDM) induced higher expression of AQP2 in the CDs and it was significantly further augmented by the addition of A83, U0126, and XAV, as shown through the reporter allele (C), protein staining (D), and qPCR (E). *n*=3 independent biological replicates per condition; column and error bars represent mean and standard deviation, respectively; P-values shown in figure. Scale bars, 500 μm (A, D), 1,000 μm (C) and 100 μm (F).

Our temporal transcriptomic profiling revealed that early stages of UB differentiation at days 3 and 7 appeared largely congruent with known developmental paradigms (Fig. 5E and Supp. Fig. 8A-C), so we focused on whether modifications of the later stages could rescue principal cell specification and maturation. In our prior study, isolated UB organoids rapidly acquired AQP2 expression when they were transitioned from a progenitor supportive medium (UB Media) to a more minimal medium devoid of growth factors^16^ (Supp. Fig. 8D-E), which is termed CD Medium and contains the hormones arginine vasopressin and aldosterone. When the recombinant organoids were similarly switched from Mix Medium to the CD Medium at day 10, they acquired faint AQP2 expression by day 14 (Fig. 7B-D), indicating that this potential was maintained by day 10 but maturation was still largely suppressed. To overcome this deficiency, we empirically screened developmental signaling pathways in isolated 3D UB organoids and found that activation of either WNT, TGFB, or FGF/RTK signaling was sufficient to repress reporter activation (Supp. Fig. 8E-F). To translate these findings to the rescue of maturation in the recombinant organoids, we supplemented the CD Medium with inhibitors of all three pathways (‘AUX’; A83-01, U0126, and XAV939). Remarkably, four-day exposure to this condition led to robust activation of the *AQP2* reporter (Fig. 7B-C) and a significant increase in expression of this and other CD markers (*ELF5*, *SCNN1B*) measured via qPCR (Fig. 7E). Wholemount staining further confirmed the induction of AQP2^+^ CDs in the organoids (Fig. 7D), and it reassuringly showed that these manipulations did not grossly impact the NM lineage, epithelial fusions, or overall organoid organization (Supp. Fig. 9A-B). Thus, this further advanced the organoid outcomes to having appropriately polarized and segmented nephrons that fuse distally to matured AQP2^+^ CDs (Fig. 7F), producing the most authentically organized kidney organoids yet described.

## Discussion

Here we established an hPSC-derived co-culture system to recapitulate essential interactions between the NM and UB in development that led to more advanced and functionally organized human kidney organoids compared to existing methodologies. In this model, induced NM undergoes a stereotyped sequence of nephrogenesis that culminates in fusion of the distal pole to the UB-derived CD, establishing for the first time nephrons that are both properly polarized and connected to CD-like structures. This robust system for establishing the luminal connection, which occurs both *in vitro* and following transplantation *in vivo*, that is required for passage of tubular fluid in the nephron into the collecting system is a key milestone toward the generation of more functional renal tissues from hPSCs. We further defined strategies for augmenting the frequency of anastomoses by applying developmental principles to extrinsically modulate the ratio of proximal-distal differentiation in the nephron segments. The induced UB epithelium similarly develops in a stepwise fashion, and we identified conditions to enhance terminal maturation of the integrated CDs within the organoids.

While the ability to produce *de novo* nephron-like structures from hPSCs has been replicated in countless studies over the past decade^6,7,36^, it previously had not been determined whether they were competent to establish a drainage mechanism via controlled epithelial fusion. In addition to its importance to forming functional tissue *in vitro*, replicating this process has also been proposed as a strategy to integrate hPSC-derived nephrons into the host collecting system *in vivo*. Our advanced organoid model described here provides a solution to this longstanding question and dilemma. The fusogenic properties of the early distal tubule are indeed recapitulated in hPSC-derived nephrons, and the developmental process occurs in a predictable manner when provided with appropriately staged UB epithelia. Although dozens of nephrons fuse to the UB in our chimeric organoids, there are many others that do not. Thus far, our analyses indicate that one essential criterion is the formation of a GATA3^+^ distal domain, which does not occur universally in all the renal vesicles. Upon observing fusion events in hundreds of organoids, we postulate that other requirements include the proper orientation and some minimum proximity of the distal domain relative to the UB. Unlike *in vivo*, where these variables are reproducible and tightly controlled through spatiotemporally conserved nephron morphogenesis, they are more stochastically regulated during *in vitro* nephrogenesis. Thus while NOTCH inhibition was sufficient to increase the number of nephrons with GATA3^+^ distal ends, future work to engineer more precision and control over nephron patterning is needed to further advance this model system and increase the overall efficiency of fusion to the CDs.

Despite its central role in the formation of functional nephrons, our mechanistic understanding of the fusion process in kidney development is exceptionally limited. In mouse genetic models, ectopic proximalization of forming nephrons through NOTCH activation prohibited fusion with the CD^4^. Using our human-based system, here we provide complementary evidence demonstrating that NOTCH inhibition was sufficient to promote distal tubules with this ability (Fig. 4F), and these data collectively substantiate a model in which proximal vs. distal specification is a key deterministic event controlling this potential. However, it is yet to be determined whether NOTCH is repressing fusion competence solely through its effect on specifying segmental identity^27–29^ or also possibly participating directly in the cell biological process. Moreover, the concomitant roles of other pathways including WNT signaling, as suggested by our ligand-receptor analyses (Fig. 5F) and prior studies^20,21^, remain to be explored. The downstream transcriptional network underlying this mechanism is also unknown. Interestingly, deletion of the essential transcription factor Hnf1b caused severe perturbations in nephrogenesis but the resulting hypoplastic tubules still exhibited fusion to the CD^37^, suggesting that only a distinct subset of the distal transcriptional program is required. Since our innovative hPSC system manifests a single wave of nephrogenesis with relatively synchronized nephron fusion, combined with the high degree of experimental and genetic tractability, it offers an unparalleled platform to address these major knowledge gaps.

The collecting system *in vivo* is an orderly, radially organized structure comprising one contiguous space and all CDs within a kidney drain into a common direction and ultimately form a single tubule, the ureter. Although the methods described here reproducibly generated organoids interlaced with extensive CD-like tubules, the pattern of growth and morphogenesis of the UB epithelia was largely stochastic and led to random CD configurations. Future efforts will be needed to engineer the UB progenitors to adopt a more consistent and organized arrangement with the goal of having a single outlet analogous to the ureter. During embryogenesis, this is achieved from a single UB that exhibits iterative branching and exponential outgrowth at one end, which is driven by interactions within the self-renewing progenitor niche. In this study, despite combining NM and UB progenitors at stages predicted to exhibit such behaviors, we did not observe maintenance of NPCs and the niche over even short periods of time (Fig. 1D). It is unclear whether this might result from failure to specify *bona fide* (FOXD1^+^) stromal progenitor cells, which have been utilized to support the niche in mouse stem cell-derived organoids^17^, or from some intrinsic deficits in the induced UB and/or NM progenitors. Further exploration of the molecular profiles and methods by which these cells are generated is required to better understand how they might be improved to enable the essential nephrogenic niche behaviors.

## Acknowledgements

We thank all members of the McCracken laboratory for their contributions to this work and Scott Rankin for his generosity and general laboratory assistance. We thank Denise Marciano, Leif Oxburgh, and Iain Drummond, as well as the larger RBK Consortium and members of the Center for Stem Cell and Organoid Medicine (CuSTOM) at CCHMC for feedback and discussions on this project. The human iPSCs (iPSC72-3 and GFP- expressing line) were generated by the CCHMC Pluripotent Stem Cell Facility (PSCF; RRID:SCR_022634), and we also thank PSCF staff for technical support. Bioinformatics support was provided by Aditi Paranjpe and the Information Services for Research facility (RRID:SCR_022622). We are also grateful to the technical expertise offered by the Bio-Imaging and Analysis Facility (RRID:SCR_022628), Integrated Pathology Research Facility (RRID:SCR_022637), and the Research Flow Cytometry Facility (RRID:SCR_022634) at CCHMC. This work was supported by funding from the Cincinnati Children’s Research Foundation (CCRF), the Pediatric Center of Excellence in Nephrology at Washington University (5P50DK133943), and a pilot grant through the (Re)Building a Kidney Consortium and ATLAS-D2K Center (U24DK135157). K.W.M. was supported by a Child Health Research Career Development Award from NIH/NICHD (K12HD028827) and a Carl W. Gottschalk Research Scholar Grant from the American Society of Nephrology. Biorender software was used for the generation of Figures 1A, 3A, 4A, 6A, 7B, and Supp. Figures 1A, 1C, 8D, and 8F.

## Author Contributions

Conceptualization, M.S. and K.W.M., Methodology, M.S., N.P.-S., N.S., M.A.H., R.K., and K.W.M., Investigation, M.S., B.C., N.P.-S., N.S., L.E., W.Z., and K.W.M., Resources, M.A.H., J.V.B., and K.W.M., Writing – Original Draft, M.S. and K.W.M., Writing – Review & Editing, M.S., N.P.-S., R.K., M.A.H., J.V.B., and K.W.M., Visualization, M.S. and K.W.M., Supervision – J.V.B. and K.W.M., Funding Acquisition, K.W.M.

## Data Availability

Single cell RNA-sequencing datasets have been deposited at the Gene Expression Omnibus (GSE271239), which will be made publicly available at the time of publication.

## Supplemental Informatio

Document S1. Supplementary Figures 1-9 and Legends.

## Methods

### Statistical analysis

No statistical methods were used to predetermine sample size. Values are presented as mean ± s.d. Replicates represent biologically independent samples. All statistical analyses were performed using Prism 8 (GraphPad Software). Statistical analyses between two groups were performed by using unpaired Student’s *t*-test if the variations were equal and unpaired Welch’s *t*-test if the variations were unequal. Statistical analyses between multiple groups (more than two groups) were performed using one-way ANOVA followed by post-hoc Tukey’s multiple comparison test. Differences with values of *P* < 0.05 were considered statistically significant. All *P* values are displayed in the figure legends. The exact N (sample size), *P* values and the statistical test used for each panel are shown in Table 1.

**Table 1.**
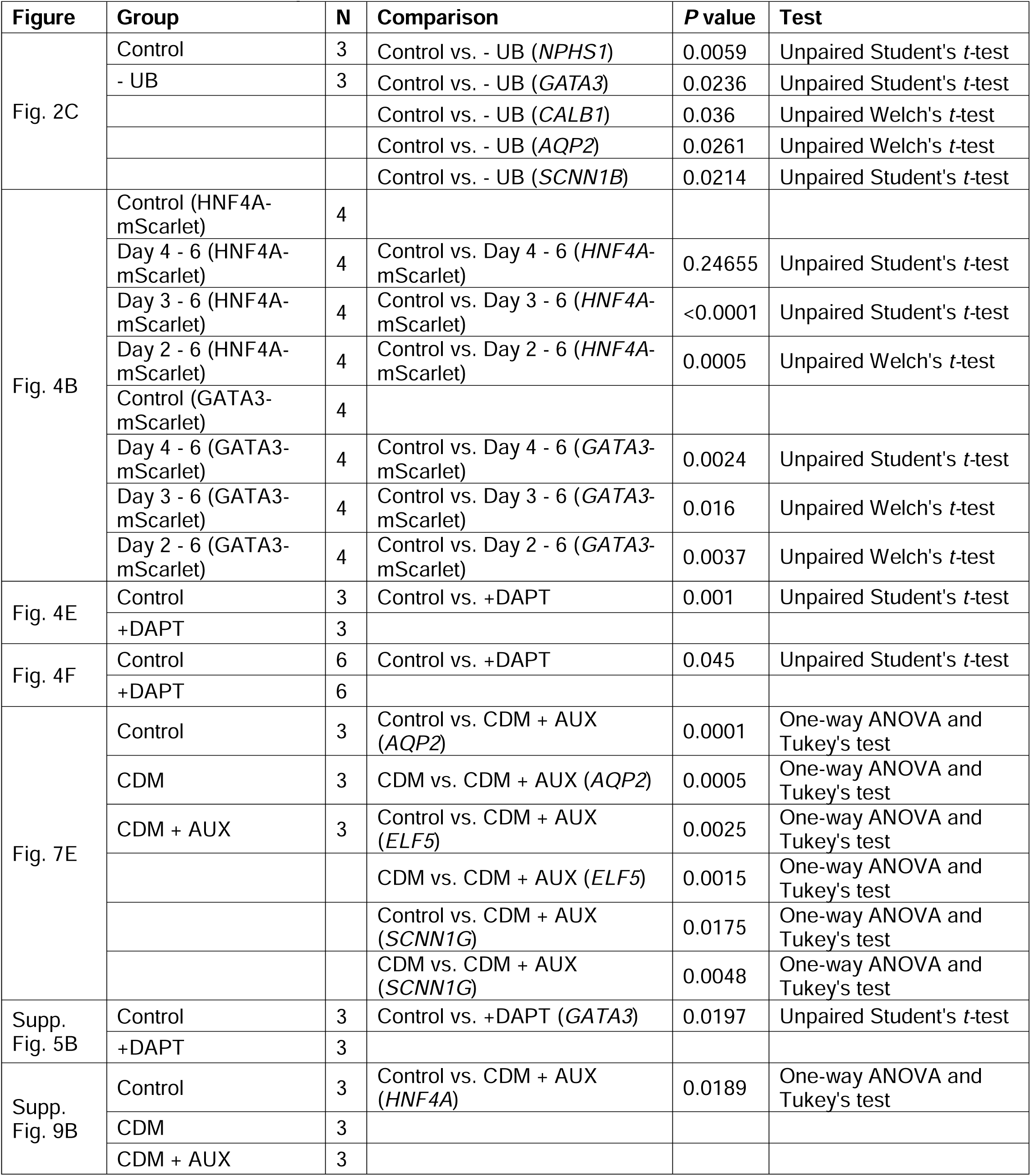
Statistical tests and p-values.

### Human PSC lines and culture

**Cell lines**. hPSC lines used in this study include H9 (WA09; obtained from WiCell), H1 (WA01; obtained from WiCell), and iPSC72-3 (generated and supplied by the Pluripotent Stem Cell Facility at Cincinnati Children’s Hospital Medical Center). The *GATA3*-mScarlet hESC line was made in H9 cells as previously described^16^, and the *HNF4A*-mScarlet reporter cell line was similarly created in H9 cells (methods below). The constitutively GFP-expressing cells used for the majority of UB differentiations were iPSC72-3 cells with a CAG-GFP construct inserted in the *AAVS1* locus^38^. The *AQP2*-mScarlet line was generated in the parental iPSC72-3 line per the methods described below.

### Maintenance culture

The hPSCs were maintained in feeder-free conditions on Cultrex Stem Cell Qualified Reduced Growth Factor Basement Membrane Extract (Bio-techne, 3434-010-02) in mTeSR1 media (STEMCELL Technologies, 05850) using six-well tissue culture plates (Falcon, 353046) in a 37°C incubator with 5% CO2. The hPSCs were routinely passaged in small colonies by Gentle Cell Dissociation Reagent (STEMCELL Technologies, 07174) at a 1:8 split ratio every 4-5 days. Studies involving hESCs were reviewed and approved by the CCHMC Institutional Biosafety Committee (IBC2022-0067) and Embryonic Stem Cell Research Oversight committee (EIP220147).

### Generation of *HNF4A*-mScarlet reporter cell line

To construct a donor template plasmid, left (981 bp) and right (961 bp) homology arms flanking the *HNF4A* stop codon were amplified by PCR (iProof, BioRad) from human genomic DNA isolated from H9 undifferentiated hESCs. The forward and reverse primers for the left homology arm were 5’-aaagcttggtaccggatccgGAAGCCATTGTTGGGATGAG-3’ and 5’-acagggagaagttagtggcgccGATAACTTCCTGCTTGGTGATG-3’, with the highlighted portion representing plasmid homologous sequence used in subsequent HIFI cloning. The left and right primers for the right homology arm were 5-attatacgaagttatgagctCAAGCCGCTGGGGCTTG-3’ and 5’- gccatggcctgcagggagctATCATCCCTCTCCCACACCA-3’. The PCR fragments were purified and cloned into pUC57 vector flanking a P2A-mScarlet and PGK-HygroR cassettes. The pX458 plasmid (Addgene 48138, kindly provided by Feng Zhang) was used to deliver Cas9 and guide RNA (gRNA) targeting the 3’ side of the *HNF4A* stop codon. To introduce the gRNA, the forward and reverse oligonucleotides (5’- CACCGAGTTATCTAGCAAGCCGCTG-3’ and 5’-aaacCAGCGGCTTGCTAGATAACTC-3’, respectively) were annealed and ligated into BbsI-digested pX458 plasmid. Plasmid sequences were verified using Sanger sequencing.

The donor template and Cas9/gRNA plasmids were reverse transfected into H9 hESCs using TransIT-LT1 (Mirus) according to manufacturer recommendations. Prior to transfection, H9 cells were dissociated with Accutase (Stem Cell Technologies) and plated into 6-well plates in mTeSR1 with ROCK inhibitor Y-27632 (10 μM; Cayman Chemical) at a concentration of 1.0 x 10^6^ cells per well. Beginning the following day, the medium was replaced with fresh mTeSR1 (without ROCK inhibitor) daily until cells were ready for passage. Two days following passage with Gentle Cell Dissociation Reagent, the cells were exposed to Hygromycin (50 μg ml^-1^) for selection. Resistant clones emerged and were identified as healthy and normal-appearing colonies that were growing in Hygromycin more than 5 days after starting selection. The individual clones were then expanded, genotyped, and tested in kidney organoid differentiation.

### Generation of *AQP2*-mScarlet reporter cell line

The donor template plasmid for AQP2 was synthesized (Genewiz) to contain left (826 bp) and right (875 bp) homology arms surrounding the endogenous stop codon that flanked the P2A-mScarlet and PGK-HygroR cassettes in the pUC57 vector. Forward and reverse oligonucleotides (5’-CACCGAGCGTCCGTCGGGGCCGTAG-3’ and 5’-aaacCTACGGCCCCGACGGACGCTC-3’, respectively) were similarly annealed and ligated into BbsI-digested pX458 to produce the Cas9/gRNA vector. Transfection, selection, and clonal expansion was performed as described above for the HNF4A cell line.

### Generation of UB spheroids from hPSCs

#### Directed differentiation protocol

hPSCs were differentiated into UBs using published methods^16,19^. In brief, cells were dissociated with Accutase (STEMCELL Technologies, 07920) and plated onto Cultrex-coated 24-well or 6-well plates in mTeSR1 with ROCK inhibitor (Y-27632; 10 μM). On the next day (day 0), cells were induced into a primitive streak-like fate in basic differentiation medium, consisting of Advanced RPMI 1640 (Thermo Scientific, 12633020) and 1x L-GlutaMAX (Thermo Scientific, 35050-061), supplemented with 5 μM CHIR99021 (CHIR, Cayman Chemical), 50 ng ml^−1^ of Activin A (PeproTech), 25 ng ml^−1^ of BMP4 (PeproTech), and 25 ng ml^−1^ of FGF2 (PeproTech). On day 1, after 25 to 30 hours depending on the rate of differentiation^19^, the media were replaced with basic differentiation medium containing 25 ng ml^−1^ FGF2, 1 μM A83-01 (Cayman Chemical), 0.1 μM LDN193189 (LDN; Cayman Chemical) and 0.1 μM RA (Sigma-Aldrich) for 2 days to induce anterior intermediate mesoderm on day 3. Media were changed daily.

On day 3, cells were dissociated with Accutase and aggregated in AggreWell-400 24-well plate (STEMCELL Technologies, 34411) in Basic Differentiation Medium supplemented with 50 ng ml^−1^ of FGF9 (R&D Systems) and 0.1 μM RA. At day 5, half-medium change was performed with medium containing 100 ng ml^−1^ GDNF (PeproTech) and 0.1 μM RA. By day 6, the UB spheroids exhibited typical morphology and were collected to mix with NM.

#### Cryopreservation of progenitor cells

To facilitate more flexible coordination of the two differentiation protocols, we developed methods to cryopreserve UB progenitor cells at day 3. The cells were dissociated with Accutase as described above and pelleted through centrifugation at 300 x *g* for 3 minutes. The supernatant was then aspirated and the cells were re-suspended in freezing medium comprising 45% Basic Differentiation Medium, 45% Fetal Bovine Serum (FBS; Thermo Scientific, 10437-028), and 10% Dimethyl Sulfoxide (DMSO; Fisher BioReagents, BP231-100). This suspension was dispensed into cryovials in 1 mL aliquots containing 1.5-3.0 x 10^6^ cells, which is enough to seed two wells of the AggreWell plates upon thawing. The vials were frozen in Mr. Frosty freezing containers with isopropyl alcohol at −80°C, and they were transferred to liquid nitrogen storage in the subsequent 1-5 days. To generate spheroids, frozen cells were rapidly thawed, centrifuged, resuspended in medium and plated into AggreWell plates as described above. To date, we have used frozen cells for >9 months with no observable difference in survival or differentiation outcomes.

#### Formation of branching 3D UB organoids

To test signaling pathways that might inhibit maturation of CD epithelial cells, we grew three-dimensional UB/CD organoids as previously described^16,19^. In brief, day 6 UB spheroids were collected and embedded into 100% Matrigel Matrix (Corning, 354234) by spotting 45-μl droplets in 24-well plates (Thermo Fisher Scientific, 142475). Typically, the spheroids (∼1,200) from one well of the AggreWell were used to generate 12-24 wells of UB organoids. The plate was placed in 37°C incubator for 60 minutes to solidify the Matrigel and then overlaid with basic differentiation medium containing 50 ng ml−1 of GDNF (PeproTech), 50 ng ml−1 of FGF10 (PeproTech), 2 μM CHIR, 0.1 μM LDN, 1 μM A83-01, 0.1 μM RA and 10 μM Y27632 (Cayman Chemical) for 7 days. The medium was changed after 3-4 days. On day 13, we switched to CD differentiation medium that comprised the same basic differentiation medium supplement with 10 nM arginine vasopressin (Sigma-Aldrich) and 10 nM aldosterone (Sigma-Aldrich) to induce CD maturation. For inhibiting maturation assay, the following growth factors were added: 50 ng ml^−1^ Activin A, 50 ng ml^−1^ BMP4, 50 ng ml^−1^ FGF7 (PeproTech), 50 ng ml^−1^ FGF10, 50 ng ml^−1^ GDNF, 3 μM CHIR and 0.1 μM RA.

### Generation of nephrogenic mesenchyme from hPSCs

#### Directed differentiation protocol

hPSCs were differentiated into NM using methods adapted from published protocols^7,39^. Briefly, cells were dissociated with Accutase (STEMCELL Technologies, 07920) and plated onto Cultrex-coated 6-well plates. Differentiation was start on the following day (day 0), with the same basal medium used at all steps. Cells were exposed to 8 μM CHIR (with 5 ng ml^−1^ Noggin if necessary) from days 0-4, followed by 10 ng ml^−1^ Activin A on days 4-7, and 10 ng ml^−1^ FGF9 from day 7-8. Media were changed daily. At day 8, the cells were dissociated with Accutase, and collected to mix in aggregates with UBs or cryopreserved in liquid nitrogen (details are shown below).

#### Cryopreservation of progenitor cells

NM progenitors at day 8 were cryopreserved using similar methods as described above for UB progenitors. Cells were dissociated with Accutase, pelleted by centrifugation, and resuspended in the same freezing medium. The cell suspension was aliquoted into cryovials at concentrations between 5-15 x 10^6^ cells ml^−1^, placed in a Mr. Frosty freezing container and stored at −80°C overnight. The next day, cryovials were moved into liquid nitrogen for long-term storage. To thaw cells, cryovials were warmed by hand until the ice was almost completely thawed. The suspension was then transferred to a 15 ml conical tube containing 5 ml DMEM (Thermo Scientific, 11965092) and centrifuged at 300 x *g* for 3 minutes. NM cell pellets were directly used to mix with UBs and form organoids after aspirating supernatant. To date, frozen NM cells have been used for >8 months with no observable loss of differentiation efficiency.

### Assembling progenitor cells into kidney organoids

To generate kidney organoids, we aggregated NM with or without UB spheroids at high density on transwell filter membranes using previously described aggregation techniques with modifications^40^. On day 0, intact day 6 UB spheroids were collected from AggreWell plate with a P1000 micropipette and transferred to a 1.7 ml Posi-Click Microcentrifuge Tube (Denville Scientific Inc., C2170). Day 8 NM progenitors (either freshly differentiated and dissociated or, more commonly, thawed cells) were collected via centrifugation and added to the same microcentrifuge tube at a ratio of 5.0 x 10^5^ NM progenitors per ∼50 UB spheroids to generate one organoid. For best aggregation results, we mixed enough NM cells (3.0-6.0 x 10^6^) and UB spheroids (300-600; corresponding to ¼ to ½ of a single AggreWell well) to make 6-12 organoids in a single tube. The mixture was centrifuged at 450 x *g* for 4 minutes, and the supernatant was carefully aspirated with a micropipette to remove as much medium as possible. The pellet was gently resuspended at 3.75 x 10^5^ NM cells μl^-1^ in basic differentiation medium and the dense suspension was spotted in 1.33 μl drops onto a transparent PET transwell insert membrane (Falcon, 353090, 0.4 μm pore size). Up to 6 organoids could be spatially arranged on each filter. Differentiation medium was added only to the lower chamber of the well to create air-liquid interface cultures. At the time of aggregation, 1.3 ml media consisting of 90% basic differentiation media, 10% FBS, and supplemented with 0.2 μM LDN193189 and 10 μM Y27632 (Cayman Chemical) was added into the lower compartment. After 5 hours, the medium was replaced with 1.3 ml medium containing 10% FBS with 0.2 μM LDN193189 but without Y27632. From day 2-14, the medium only contained basic differentiation medium with 10% FBS. Media were changed daily from days 0-4 (with 1.3 ml) and every 2 days (with 1.6 ml) afterward. For NM-only organoids without UB, the same protocol was used but with 6.0 x 10^5^ cells/organoid to account for the estimated cell number in the UB spheroids. For UB-only air-liquid interface cultures, day 6 UB spheroids were pelleted, resuspended at ∼200 spheroids/μl in basic differentiation medium and spotted at 1.5 μl each onto the Transparent PET Membrane.

For the Notch inhibition experiments, 10 μM DAPT (Cayman Chemical) was added to the above culture medium for varying lengths of time between days 2-6. For improving the terminal maturation of CD epithelia assay, at day 10 the organoid medium was changed to basic differentiation medium supplemented with 10 nM arginine vasopressin (Sigma-Aldrich), 10 nM aldosterone (Sigma-Aldrich), 3 μM A83-01, 5 μM U0126 (Cayman Chemical) and 1 μM XAV939 (Cayman Chemical). The organoids were then cultured to day 14 for analysis.

### RNA isolation and qRT–PCR

Total RNA was isolated using NucleoSpin RNA Plus kit (Macherey-Nagel, 740984), and reverse-transcribed using iScript cDNA synthesis kit (Bio-Rad, 1708841). qRT-PCR was performed on QuantStudio 3 Real-Time PCR System (Thermo Fisher Scientific) using iTaq Universal SYBR Green Supermix (Bio-Rad, 1725124). Relative mRNA expression levels were normalized to GAPDH or PPIA gene expression by the ΔΔCT method. Primer sequences are listed in Table 2.

**Table 2.**
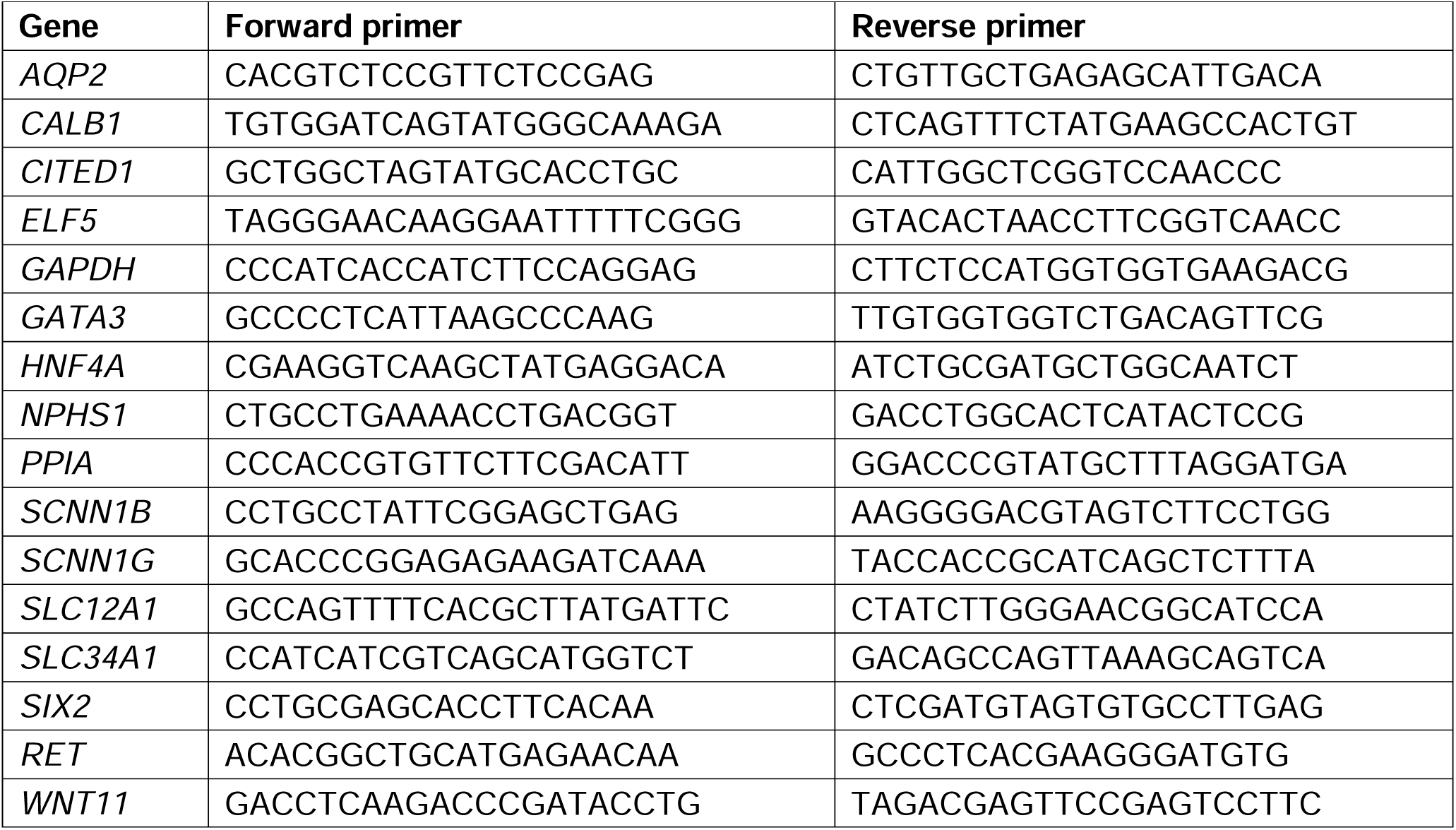
qPCR Primers.

### Immunofluorescent staining and histology

For whole-mount staining, cultured organoids were directly fixed on the transwell in 4% paraformaldehyde (in PBS) for 1 hour at room temperature, and transplanted organoids were dissected from under the renal capsule and fixed in 4% paraformaldehyde (in PBS) for 2 hours at room temperature. Following fixation, organoids were washed thoroughly in PBS for three times for five minutes. For staining, the organoids were incubated in blocking buffer (0.5% Triton X-100 and 5% normal donkey serum in PBS) for 1 hour at room temperature and incubated with primary antibodies overnight at 4°C in blocking buffer. The following day, the organoids were washed with PBS three times and incubated with secondary antibodies and DAPI (Sigma-Aldrich) for 4 hours at room temperature. After washing three times with PBS, organoids from the transwell were transferred to glass slides, mounted with Fluoromount G (Invitrogen), coverslipped and imaged by confocal microscopy (Nikon A1R inverted confocal microscope). For transplanted organoids, which were larger, imaging was performed directly in PBS without mounting. Primary and secondary antibodies used are listed in Table 3.

**Table 3.** Primary and Secondary Antibodies.

| Target | Vendor | Catalog Number | Dilution |
| --- | --- | --- | --- |
| AQP2 | Abcam | ab199975 | 1:1000 |
| CALB1 | Sigma-Aldrich | C9848 | 1:200 |
| CDH1 | BD Biosciences | 610181 | 1:500 |
| CDH1 | R&D systems | AF648 | 1:1000 |
| GATA3 | Abcam | ab199428 | 1:300 |
| GATA3 | Cell Signaling Technologies | 5852S | 1:300 |
| HNF4A | Santa Cruz Biotechnology | sc-374229 | 1:100 |
| HNF4A | Cell Signaling Technologies | 3113S | 1:200 |
| JAGGED1 | R&D systems | AF599 | 1:500 |
| Laminin | Sigma-Aldrich | L9393 | 1:200 |
| LHX1 | Abcam | ab229474 | 1:200 |
| NPHS1 | R&D systems | AF4269 | 1:500 |
| PAX2 | R&D systems | AF3364 | 1:500 |
| PDGFRB | R&D systems | AF385 | 1:200 |
| PECAM1 | Cell marque | 131R-24 | 1:200 |
| POU3F3 | Novus Biologicals | NBP1-49872 | 1:200 |
| PRKCZ | Santa Cruz Biotechnology | sc-17781 | 1:200 |
| RET | R&D systems | AF1485 | 1:500 |
| SIX1 | Cell Signaling Technologies | 12891 | 1:500 |
| SIX2 | Proteintech | 11562-1-AP | 1:200 |
| SLC12A1 | Invitrogen | PA5-80003 | 1:200 |
| WT1 | Boster Bio | M00199-1 | 1:250 |
| ZO-1 | Invitrogen | 33-9100 | 1:200 |
| **Secondary Antibodies** |  |  |  |
| Alexa Fluor 488 Donkey Anti-Mouse IgG | Jackson ImmunoResearch | 715-545-150 | 1:500 |
| Alexa Fluor 488 Donkey Anti-Rabbit IgG | Jackson ImmunoResearch | 711-545-152 | 1:500 |
| Alexa Fluor 488 Donkey Anti-Goat IgG | Jackson ImmunoResearch | 705-545-147 | 1:500 |
| Alexa Fluor 594 Donkey Anti-Mouse IgG | Jackson ImmunoResearch | 715-585-150 | 1:500 |
| Alexa Fluor 594 Donkey Anti-Rabbit IgG | Jackson ImmunoResearch | 711-585-152 | 1:500 |
| Alexa Fluor 594 Donkey Anti-Goat IgG | Jackson ImmunoResearch | 705-585-147 | 1:500 |
| Alexa Fluor 647 Donkey Anti-Mouse IgG | Jackson ImmunoResearch | 715-605-150 | 1:500 |
| Alexa Fluor 647 Donkey Anti-Rabbit IgG | Jackson ImmunoResearch | 711-605-152 | 1:500 |
| Alexa Fluor 647 Donkey Anti-Goat IgG | Jackson ImmunoResearch | 705-605-147 | 1:500 |

For frozen sectioning, transplanted tissues were fixed in 4% paraformaldehyde, washed by PBS thoroughly and incubated with 25% sucrose (in PBS) overnight at 4 °C. Then the samples were mounted in OCT compound (Thermo Fisher Scientific), frozen in blocks overnight, cut into 7-μm sections by cryostat and placed on slides. For staining, sections were incubated in blocking buffer (0.1% Triton X-100 and 5% normal donkey serum in PBS) for 1 hour at room temperature and incubated with primary antibodies in blocking buffer overnight at 4 °C. Then slides were washed by PBS, incubated with secondary antibodies and DAPI for 1 hour at room temperature and washed by PBS. Slides were mounted with Fluoromount G and coverslipped.

For histology, implanted organoids were fixed in 4% paraformaldehyde at room temperature and subjected to overnight processing followed by paraffin embedding. The blocks were cut into 5 μm sections by a microtome. Hematoxylin and Eosin (H&E) staining was performed following the manufacturer’s directions. Histological images were captured by widefield microscopy (Nikon 90i upright widefield microscope).

### Image analysis and quantification

To examine and quantify nephron-UB fusion events, we examined 3D projections of confocal z-stacks of wholemount-stained organoids using Imaris. All continuous luminal connections were identified and quantified manually using TJP1, GFP, and Dapi staining in day 14 organoids. We further quantified the number of fused epithelia that expressed GATA3 on the GFP^-^ end of the junction. To adjust for the varying density of UB-derived CDs in the organoids, the frequency of connections was normalized to the GFP^+^ area (mm^2^), as measured in Imaris software. To quantify proximal and distal specification in the DAPT-treated organoids, we used organoids generated from *HNF4A*-mScarlet and *GATA3*-mScarlet cell lines, respectively. The percentage of reporter-positive area within the organoid was measured using General Analysis methods in Nikon NIS Elements software.

### Flow cytometry

Organoids were dissociated with TrypLE Express Enzyme (Thermo Scientific, 12605010) for 12 minutes at 37°C, followed by gentle pipetting. To fully dissociate tissue into single cells, organoids was placed back into 37 °C for incubating 3-5 more minutes and pipetted gently. Then the cells were pelleted and incubated with 200 μl LIVE/DEAD™ Fixable Blue Stain buffer (dilution 1:1000 in PBS; Invitrogen, L23105) for staining 30 min on ice. After LIVE/DEAD staining, cells were washed once with cold PBS and fixed in 1% paraformaldehyde for 1 hour on ice. We then transferred cells into a Polystyrene Test Tube (Falcon, 352235) though the cell strainer snap cap and performed analysis using a flow cytometry (LSR Fortessa, BD Biosciences). Organoids without mScarlet and GFP reporters were used as negative controls to establish gating parameters. Data were analyzed using FlowJo software.

### *In vivo* transplantation of kidney organoids

Kidney organoids were transplanted beneath the kidney capsule of NSG (NOD scid gamma) mice. The NSG mouse colony was housed and maintained in the vivarium at Cincinnati Children’s Hospital Medical Center (CCHMC). The facility is on a 14-hour/10-hour light/dark cycle and maintained at a temperature of 22°C. The veterinary facilities at CCHMC are accredited by AAALAC (000492), and all animal experiments were approved by the Institutional Animal Care and Use Committee (IACUC2021-0060 and IACUC2021-0054).

All mice used for transplantation were male and between 8-16 weeks old. Kidney organoids on day 3 following integration of NM and UB spheroids were manually removed from the transwell and transplanted into the left renal subcapsular space as described previously^35,41^. Briefly, the mouse was anesthetized using 2% inhaled isoflurane (Butler Schein). The left flank was prepared and cleansed with isopropyl alcohol and povidone-iodine, and a 1 cm vertical incision was made in the left paraspinal area. The left kidney was exposed through the incision, and the capsule was gently dissected with a probe and forceps to create a subcapsular space or pocket to hold the organoid. Two organoids were then inserted to fit snugly in the pocket, and the kidney was returned to the retroperitoneal space. An intraperitoneal injection of piperacillin-tazobactam (100 mg kg^-1^) was administered for antimicrobial prophylaxis, and the surgical incision was sutured closed. Postoperatively, mice received subcutaneous Buprenex (0.05 mg kg^-1^) for analgesia. Transplanted mice were sacrificed using CO_2_ two weeks later, and the kidneys were harvested for fixing, histology, and immunofluorescent staining.

### Single cell capture, library preparation, and sequencing

For the scRNA-seq experiment, recombined organoids from days 3, 7 and 15 were dissociated by incubating with TrypLE Express for 15-17 minutes at 37 °C with intermittent trituration using a P1000 pipette until the organoids were largely dissociated into single cells. DMEM was added at a 2:1 volume ratio to the cell suspension, which was then mixed, transferred to a 15 ml conical tube, and centrifuged at 300 x *g* for 3 minutes. The cell pellet was resuspended in PBS with 0.04% bovine serum albumin (BSA; Thermo Fisher Scientific), filtered through the cell strainer snap cap tube, and transferred to a 1.7 ml Posi-Click Microcentrifuge Tube. Hashtags were then added to samples using Cell Multiplexing Oligos from 3’ CellPlex Kit Set A (10X Genomics, 1000261), according to manufacturer’s recommendations. Briefly, the single cell suspension was pelleted by centrifugation, resuspended in 100 μl multiplexing oligo, and incubated for 5 minutes at room temperature. After incubation, cells were moved to ice, washed with ice-cold wash buffer (PBS + 10% FBS), and re-pelleted by centrifugation. These washes were repeated for a total of four times. All centrifugation was done at 400 x *g* at 4°C. After washes, the cells were resuspended in wash buffer at a concentration of 1,500 cells/μl. The cell suspensions were then directly delivered to the Single Cell Genomics Facility at CCHMC for quality control and library preparation. Single cell libraries were created from the pooled and multiplexed samples using the Next GEM Single Cell 3’ Assay (10X Genomics; v3.1), and sequenced on a NovaSeq 6000 instrument.

### scRNA-seq data analysis

Raw sequencing data were processed and de-multiplexed using Cell Ranger v7.0.1 with the cellranger multi pipeline. The sequence of the GFP transgene was appended to the hg38 reference genome for the alignment. Overall, sequencing yielded an average depth of 42,000 reads per cell, with an average of 18,334 UMI per cell and 5,518 genes per cell. Data were further processed in the R package Seurat (v5.0.3), where the datasets from days 3, 7, and 15 were merged and then analyzed using a standard pipeline for normalization, scaling, cell cycle regression, and dimensional reduction. Cells expressing less than 500 genes or with >20% of reads mapping to mitochondrial genes were excluded from the final dataset. The top 2,000 variable genes were used with a dimension value of 18 for dimensional reduction and clustering via Uniform Manifold Approximation and Projection (UMAP). Figures and data visualization were generated using Seurat and the package scCustomize (v2.1.2). To identify candidate ligand-receptor interactions, data were analyzed using CellChat^42^ (v1.5.0).

## References

1. Mugford, J.W., Sipila, P., McMahon, J.A., and McMahon, A.P. (2008). Osr1 expression demarcates a multi-potent population of intermediate mesoderm that undergoes progressive restriction to an Osr1-dependent nephron progenitor compartment within the mammalian kidney. Dev Biol 324, 88–98. 10.1016/j.ydbio.2008.09.010.

2. Taguchi, A., Kaku, Y., Ohmori, T., Sharmin, S., Ogawa, M., Sasaki, H., and Nishinakamura, R. (2014). Redefining the in vivo origin of metanephric nephron progenitors enables generation of complex kidney structures from pluripotent stem cells. Cell Stem Cell 14, 53–67. 10.1016/j.stem.2013.11.010.

3. McMahon, A.P. (2016). Development of the Mammalian Kidney. Curr Top Dev Biol 117, 31–64. 10.1016/bs.ctdb.2015.10.010.

4. Kao, R.M., Vasilyev, A., Miyawaki, A., Drummond, I.A., and McMahon, A.P. (2012). Invasion of distal nephron precursors associates with tubular interconnection during nephrogenesis. J Am Soc Nephrol 23, 1682–1690. 10.1681/ASN.2012030283.

5. Georgas, K., Rumballe, B., Valerius, M.T., Chiu, H.S., Thiagarajan, R.D., Lesieur, E., Aronow, B.J., Brunskill, E.W., Combes, A.N., Tang, D., et al. (2009). Analysis of early nephron patterning reveals a role for distal RV proliferation in fusion to the ureteric tip via a cap mesenchyme-derived connecting segment. Dev Biol 332, 273–286. 10.1016/j.ydbio.2009.05.578.

6. Takasato, M., Er, P.X., Chiu, H.S., Maier, B., Baillie, G.J., Ferguson, C., Parton, R.G., Wolvetang, E.J., Roost, M.S., Chuva de Sousa Lopes, S.M., and Little, M.H. (2015). Kidney organoids from human iPS cells contain multiple lineages and model human nephrogenesis. Nature 526, 564–568. 10.1038/nature15695.

7. Morizane, R., Lam, A.Q., Freedman, B.S., Kishi, S., Valerius, M.T., and Bonventre, J.V. (2015). Nephron organoids derived from human pluripotent stem cells model kidney development and injury. Nat Biotechnol 33, 1193–1200. 10.1038/nbt.3392.

8. Lawlor, K.T., Vanslambrouck, J.M., Higgins, J.W., Chambon, A., Bishard, K., Arndt, D., Er, P.X., Wilson, S.B., Howden, S.E., Tan, K.S., et al. (2020). Cellular extrusion bioprinting improves kidney organoid reproducibility and conformation. Nat Mater. 10.1038/s41563-020-00853-9.

9. Homan, K.A., Gupta, N., Kroll, K.T., Kolesky, D.B., Skylar-Scott, M., Miyoshi, T., Mau, D., Valerius, M.T., Ferrante, T., Bonventre, J.V., et al. (2019). Flow-enhanced vascularization and maturation of kidney organoids in vitro. Nat Methods 16, 255–262. 10.1038/s41592-019-0325-y.

10. Vanslambrouck, J.M., Wilson, S.B., Tan, K.S., Groenewegen, E., Rudraraju, R., Neil, J., Lawlor, K.T., Mah, S., Scurr, M., Howden, S.E., et al. (2022). Enhanced metanephric specification to functional proximal tubule enables toxicity screening and infectious disease modelling in kidney organoids. Nat Commun 13, 5943. 10.1038/s41467-022-33623-z.

11. Grote, D., Souabni, A., Busslinger, M., and Bouchard, M. (2006). Pax 2/8-regulated Gata 3 expression is necessary for morphogenesis and guidance of the nephric duct in the developing kidney. Development 133, 53–61. 10.1242/dev.02184.

12. Uchimura, K., Wu, H., Yoshimura, Y., and Humphreys, B.D. (2020). Human Pluripotent Stem Cell-Derived Kidney Organoids with Improved Collecting Duct Maturation and Injury Modeling. Cell Rep 33, 108514. 10.1016/j.celrep.2020.108514.

13. Howden, S.E., Wilson, S.B., Groenewegen, E., Starks, L., Forbes, T.A., Tan, K.S., Vanslambrouck, J.M., Holloway, E.M., Chen, Y.H., Jain, S., et al. (2021). Plasticity of distal nephron epithelia from human kidney organoids enables the induction of ureteric tip and stalk. Cell Stem Cell 28, 671–684 e676. 10.1016/j.stem.2020.12.001.

14. Taguchi, A., and Nishinakamura, R. (2017). Higher-Order Kidney Organogenesis from Pluripotent Stem Cells. Cell Stem Cell 21, 730–746 e736. 10.1016/j.stem.2017.10.011.

15. Mae, S.I., Ryosaka, M., Toyoda, T., Matsuse, K., Oshima, Y., Tsujimoto, H., Okumura, S., Shibasaki, A., and Osafune, K. (2018). Generation of branching ureteric bud tissues from human pluripotent stem cells. Biochem Biophys Res Commun 495, 954–961. 10.1016/j.bbrc.2017.11.105.

16. Shi, M., McCracken, K.W., Patel, A.B., Zhang, W., Ester, L., Valerius, M.T., and Bonventre, J.V. (2023). Human ureteric bud organoids recapitulate branching morphogenesis and differentiate into functional collecting duct cell types. Nat Biotechnol 41, 252–261. 10.1038/s41587-022-01429-5.

17. Tanigawa, S., Tanaka, E., Miike, K., Ohmori, T., Inoue, D., Cai, C.L., Taguchi, A., Kobayashi, A., and Nishinakamura, R. (2022). Generation of the organotypic kidney structure by integrating pluripotent stem cell-derived renal stroma. Nat Commun 13, 611. 10.1038/s41467-022-28226-7.

18. Tsujimoto, H., Kasahara, T., Sueta, S.I., Araoka, T., Sakamoto, S., Okada, C., Mae, S.I., Nakajima, T., Okamoto, N., Taura, D., et al. (2020). A Modular Differentiation System Maps Multiple Human Kidney Lineages from Pluripotent Stem Cells. Cell Rep 31, 107476. 10.1016/j.celrep.2020.03.040.

19. Shi, M., Fu, P., Bonventre, J.V., and McCracken, K.W. (2023). Directed differentiation of ureteric bud and collecting duct organoids from human pluripotent stem cells. Nat Protoc 18, 2485–2508. 10.1038/s41596-023-00847-2.

20. Lindstrom, N.O., Lawrence, M.L., Burn, S.F., Johansson, J.A., Bakker, E.R., Ridgway, R.A., Chang, C.H., Karolak, M.J., Oxburgh, L., Headon, D.J., et al. (2015). Integrated beta-catenin, BMP, PTEN, and Notch signalling patterns the nephron. Elife 3, e04000. 10.7554/eLife.04000.

21. Low, J.H., Li, P., Chew, E.G.Y., Zhou, B., Suzuki, K., Zhang, T., Lian, M.M., Liu, M., Aizawa, E., Rodriguez Esteban, C., et al. (2019). Generation of Human PSC-Derived Kidney Organoids with Patterned Nephron Segments and a De Novo Vascular Network. Cell Stem Cell 25, 373–387 e379. 10.1016/j.stem.2019.06.009.

22. Howden, S.E., Vanslambrouck, J.M., Wilson, S.B., Tan, K.S., and Little, M.H. (2019). Reporter-based fate mapping in human kidney organoids confirms nephron lineage relationships and reveals synchronous nephron formation. EMBO Rep 20. 10.15252/embr.201847483.

23. Barak, H., Huh, S.H., Chen, S., Jeanpierre, C., Martinovic, J., Parisot, M., Bole-Feysot, C., Nitschke, P., Salomon, R., Antignac, C., et al. (2012). FGF9 and FGF20 maintain the stemness of nephron progenitors in mice and man. Dev Cell 22, 1191–1207. 10.1016/j.devcel.2012.04.018.

24. Shakya, R., Watanabe, T., and Costantini, F. (2005). The role of GDNF/Ret signaling in ureteric bud cell fate and branching morphogenesis. Dev Cell 8, 65–74. 10.1016/j.devcel.2004.11.008.

25. Chen, L., Clark, J.Z., Nelson, J.W., Kaissling, B., Ellison, D.H., and Knepper, M.A. (2019). Renal-Tubule Epithelial Cell Nomenclature for Single-Cell RNA-Sequencing Studies. J Am Soc Nephrol 30, 1358–1364. 10.1681/ASN.2019040415.

26. Lindstrom, N.O., Tran, T., Guo, J., Rutledge, E., Parvez, R.K., Thornton, M.E., Grubbs, B., McMahon, J.A., and McMahon, A.P. (2018). Conserved and Divergent Molecular and Anatomic Features of Human and Mouse Nephron Patterning. J Am Soc Nephrol 29, 825–840. 10.1681/ASN.2017091036.

27. Cheng, H.T., and Kopan, R. (2005). The role of Notch signaling in specification of podocyte and proximal tubules within the developing mouse kidney. Kidney Int 68, 1951–1952. 10.1111/j.1523-1755.2005.00627.x.

28. Cheng, H.T., Miner, J.H., Lin, M., Tansey, M.G., Roth, K., and Kopan, R. (2003). Gamma-secretase activity is dispensable for mesenchyme-to-epithelium transition but required for podocyte and proximal tubule formation in developing mouse kidney. Development 130, 5031–5042. 10.1242/dev.00697.

29. Duvall, K., Crist, L., Perl, A.J., Pode Shakked, N., Chaturvedi, P., and Kopan, R. (2022). Revisiting the role of Notch in nephron segmentation confirms a role for proximal fate selection during mouse and human nephrogenesis. Development 149. 10.1242/dev.200446.

30. Wilson, S.B., Howden, S.E., Vanslambrouck, J.M., Dorison, A., Alquicira-Hernandez, J., Powell, J.E., and Little, M.H. (2022). DevKidCC allows for robust classification and direct comparisons of kidney organoid datasets. Genome Med 14, 19. 10.1186/s13073-022-01023-z.

31. Yu, J., Carroll, T.J., Rajagopal, J., Kobayashi, A., Ren, Q., and McMahon, A.P. (2009). A Wnt7b-dependent pathway regulates the orientation of epithelial cell division and establishes the cortico-medullary axis of the mammalian kidney. Development 136, 161–171. 10.1242/dev.022087.

32. Carroll, T.J., Park, J.S., Hayashi, S., Majumdar, A., and McMahon, A.P. (2005). Wnt9b plays a central role in the regulation of mesenchymal to epithelial transitions underlying organogenesis of the mammalian urogenital system. Dev Cell 9, 283–292. 10.1016/j.devcel.2005.05.016.

33. Tanigawa, S., Wang, H., Yang, Y., Sharma, N., Tarasova, N., Ajima, R., Yamaguchi, T.P., Rodriguez, L.G., and Perantoni, A.O. (2011). Wnt4 induces nephronic tubules in metanephric mesenchyme by a non-canonical mechanism. Dev Biol 352, 58–69. 10.1016/j.ydbio.2011.01.012.

34. van den Berg, C.W., Ritsma, L., Avramut, M.C., Wiersma, L.E., van den Berg, B.M., Leuning, D.G., Lievers, E., Koning, M., Vanslambrouck, J.M., Koster, A.J., et al. (2018). Renal Subcapsular Transplantation of PSC-Derived Kidney Organoids Induces Neo-vasculogenesis and Significant Glomerular and Tubular Maturation In Vivo. Stem Cell Reports 10, 751–765. 10.1016/j.stemcr.2018.01.041.

35. Pode-Shakked, N., Slack, M., Sundaram, N., Schreiber, R., McCracken, K.W., Dekel, B., Helmrath, M., and Kopan, R. (2023). RAAS-deficient organoids indicate delayed angiogenesis as a possible cause for autosomal recessive renal tubular dysgenesis. Nat Commun 14, 8159. 10.1038/s41467-023-43795-x.

36. Dorison, A., Forbes, T.A., and Little, M.H. (2022). What can we learn from kidney organoids? Kidney Int 102, 1013–1029. 10.1016/j.kint.2022.06.032.

37. Heliot, C., Desgrange, A., Buisson, I., Prunskaite-Hyyrylainen, R., Shan, J., Vainio, S., Umbhauer, M., and Cereghini, S. (2013). HNF1B controls proximal-intermediate nephron segment identity in vertebrates by regulating Notch signalling components and Irx1/2. Development 140, 873–885. 10.1242/dev.086538.

38. Eicher, A.K., Kechele, D.O., Sundaram, N., Berns, H.M., Poling, H.M., Haines, L.E., Sanchez, J.G., Kishimoto, K., Krishnamurthy, M., Han, L., et al. (2022). Functional human gastrointestinal organoids can be engineered from three primary germ layers derived separately from pluripotent stem cells. Cell Stem Cell 29, 36–51 e36. 10.1016/j.stem.2021.10.010.

39. Morizane, R., and Bonventre, J.V. (2017). Generation of nephron progenitor cells and kidney organoids from human pluripotent stem cells. Nat Protoc 12, 195–207. 10.1038/nprot.2016.170.

40. Gupta, A.K., Ivancic, D.Z., Naved, B.A., Wertheim, J.A., and Oxburgh, L. (2021). An efficient method to generate kidney organoids at the air-liquid interface. J Biol Methods 8, e150. 10.14440/jbm.2021.357.

41. Watson, C.L., Mahe, M.M., Munera, J., Howell, J.C., Sundaram, N., Poling, H.M., Schweitzer, J.I., Vallance, J.E., Mayhew, C.N., Sun, Y., et al. (2014). An in vivo model of human small intestine using pluripotent stem cells. Nat Med 20, 1310–1314. 10.1038/nm.3737.

42. Jin, S., Guerrero-Juarez, C.F., Zhang, L., Chang, I., Ramos, R., Kuan, C.H., Myung, P., Plikus, M.V., and Nie, Q. (2021). Inference and analysis of cell-cell communication using CellChat. Nat Commun 12, 1088. 10.1038/s41467-021-21246-9.

