## Supplemental Figures for "Integrating collecting systems in kidney organoids through fusion of distal nephron to ureteric bud"

**Supplementary Figure Legends**

**Supplementary Figure 1. Differentiation strategies for generating NM and UB progenitor cells. A.** NM was induced in monolayer format and used to make organoids at day 8. **B.** The cultures expressed key nephron progenitor markers including SIX2, SIX1, and PAX2. **C.** UB spheroids were generated through aggregation of pronephric intermediate mesoderm progenitors at day 3 of differentiation, and the spheroids were used to assemble organoids at day 6. **D.** The spheroids comprised both UB and stromal progenitor populations as indicated in scRNA-seq analysis, with the former exhibiting high expression of the tip markers *RET*, *WNT11*, and *ETV4/5*. Scale bar, 200 μm (B).

**Supplementary Figure 2. Optimization of growth conditions for recombinant kidney organoids. A.** Addition of ROCK inhibitor Y-27632 for the first 5 hours post-mixing promoted efficient aggregation of progenitor cells and more consistent induction and epithelialization of the NM by day 4. **B.** Transient BMP inhibition (with LDN) between days 0-2 led to improved efficiency of NM induction and UB growth. **C.** Organoids generated from UBs harboring a *GATA3*-mScarlet reporter allele mixed with unlabeled (H1-derived) NM enabled visualization of the growth of the UB epithelia without seeing the UB spheroid-derived stroma. **D.** Whole-mount staining of organoids at days 0, 2, and 4 indicated that the presence of UB progenitors did not affect the rapid induction of JAG1 or the gradual extinction of SIX1 expression. **E.** Neither addition of FGF2 nor GDNF altered renal vesicle formation or UB branching by day 4, and they did not affect the differentiation of organoids by day 14. Scale bars, 500 μm (A, C), 1,000 μm (B, E) and 200 μm (D).

**Supplementary Figure 3. Luminal connection between NM-derived distal tubules and UB-derived CDs. A.** Wholemount staining demonstrated the continuous luminal connection across the junction of the $GFP^+$ CD and $GFP^-$ nephron tubule. **B.** $HNF4A^+$ proximal tubules represented the majority of NM-derived epithelial tissue in the organoids, while $GATA3^+$ ($GFP^-$) distal nephrons were far less abundant at day 14. **C.** IF staining for GATA3 and TJP1 revealed that more than one $GATA3^+$ distal tubules connected into a single $GFP^+$ ureteric tubule by day 7. Scale bars, 100 μm (A, C) and 1,000 μm (B).

**Supplementary Figure 4. Generation of reporter cell lines for *HNF4A* and *AQP2*.** Schematized strategies for CRISPR/Cas9-mediated knock-in of mScarlet reporter alleles in the human *HNF4A* (**A**) and *AQP2* (**B**) loci.

**Supplementary Figure 5. NOTCH inhibition enhances distal nephron development and fusion. A.** Exposure to DAPT from days 2-6 or 3-6 led to expansion of *GATA3* expressing tubules and nearly complete repression of podocytes (NPHS1) by day 14, whereas treatment from days 4-6 increased the distal nephron specification but maintained similar levels of proximal structures such as podocytes. **B.** Podocyte (*NPHS1*) and thick ascending limb (*SLC12A1*) were expressed at similar levels at day 14 in DAPT (d4-6)-treated and control organoids, but the distal segment marker *GATA3* was significantly increased. $n$=3 independent biological replicates per condition; column and error bars represent mean and standard deviation, respectively; $^*P$ = 0.0197. **C-D.** The $GATA3^+$ segments in DAPT-treated organoids formed normal patent anastomoses with the UB-derived CDs, and they frequently expressed the connecting segment marker CALB1. Scale bars, 1,000 μm (A) and 200 μm (C-D).

**Supplementary Figure 6. Single cell profiling of kidney organoid development. A-B.** UMAP embedding identified 16 cell clusters spanning days 3, 7 and 15 of differentiation. **C-D**. The organoids comprised nephron ($PAX2^+$/$GATA3^-$), ureteric ($PAX2^+$/$GATA3^+$), stromal (*PDGFRA*), and endothelial (*CDH5*) lineages. **E**. Dot plots of representative markers of the four different lineages in the organoids. **F.** GFP was highly expressed in nearly all cells of the ureteric clusters and a portion of the stromal lineage, but not in the nephron clusters. **G.** Clusters representing stromal (1, 4, 5, 7, 10, 13, 14) and endothelial (15) cells were identified and removed to generate the dataset shown in Figure 5A.

**Supplementary Figure 7. Lineage marker expression in kidney organoids. A-B.** Expression of representative marker genes identifying the various stages and segments of nephron and UB differentiation in the kidney organoids. **C.** EPHB and extracellular matrix signaling components were identified by CellChat analysis to be enriched in early distal nephron development compared to proximal.

**Supplementary Figure 8. Interrogating CD maturation in UB-derived epithelia. A.** UMAP embedding of UB-derived cells in the combined organoids (‘UB-Early’ and ‘UB-Late’ clusters from dataset in Figure 5A) demonstrated differentiation from tip-like progenitor cells (cluster 0) to differentiated CD cells (cluster 4). **B-C.** Despite the expression of principal cell markers, such as *ELF5* and *SCNN1B*, in cluster 4 cells, very low levels of *AQP2* were detected in the dataset. **D**. Schematic representation of methods for growing UB organoids in isolation in 3D culture and their differentiation to $AQP2^+$ cells following exposure to a minimal ‘CD Medium’. **E-F.** Following transition to CD Medium, activation of either the WNT (CHIR99021), FGF/GDNF, or TGFβ (Activin A) pathways was sufficient to repress activation of the *AQP2* reporter allele, while RA and BMP4 had no effect. Scale bar, 200 μm (E).

**Supplementary Figure 9. Organoid morphology and differentiation under conditions for CD maturation. A.** The overall morphology and architecture of organoids at day 14 was unaffected by transition to CD Medium (CDM) or CDM + AUX (A83, U0126, XAV939) culture medium, and the formation of UB-derived CD-like tubules and nephron fusion via $GATA3^+$ segments was preserved. **B.** Markers of nephron segment differentiation, including *NPHS1* (podocyte), *HNF4A* (proximal tubule), and *SLC12A1* (thick ascending limb), were largely unaffected. $n$=3 independent biological replicates per condition; column and error bars represent mean and standard deviation, respectively; *$P$ = 0.0189. Scale bar, 1,000 μm (A).

**Supplementary Figure 1**

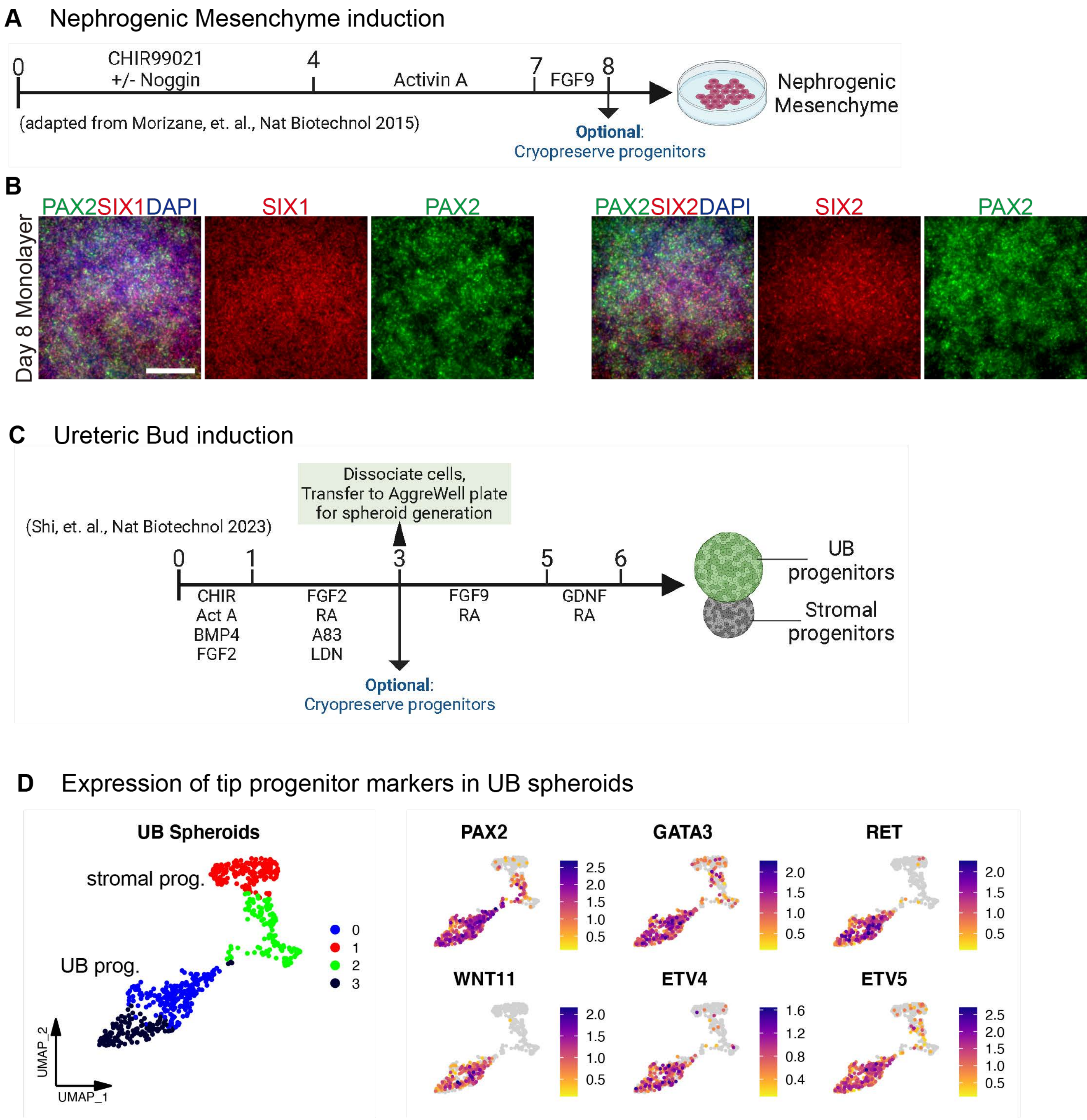

#### Supplementary Figure 2

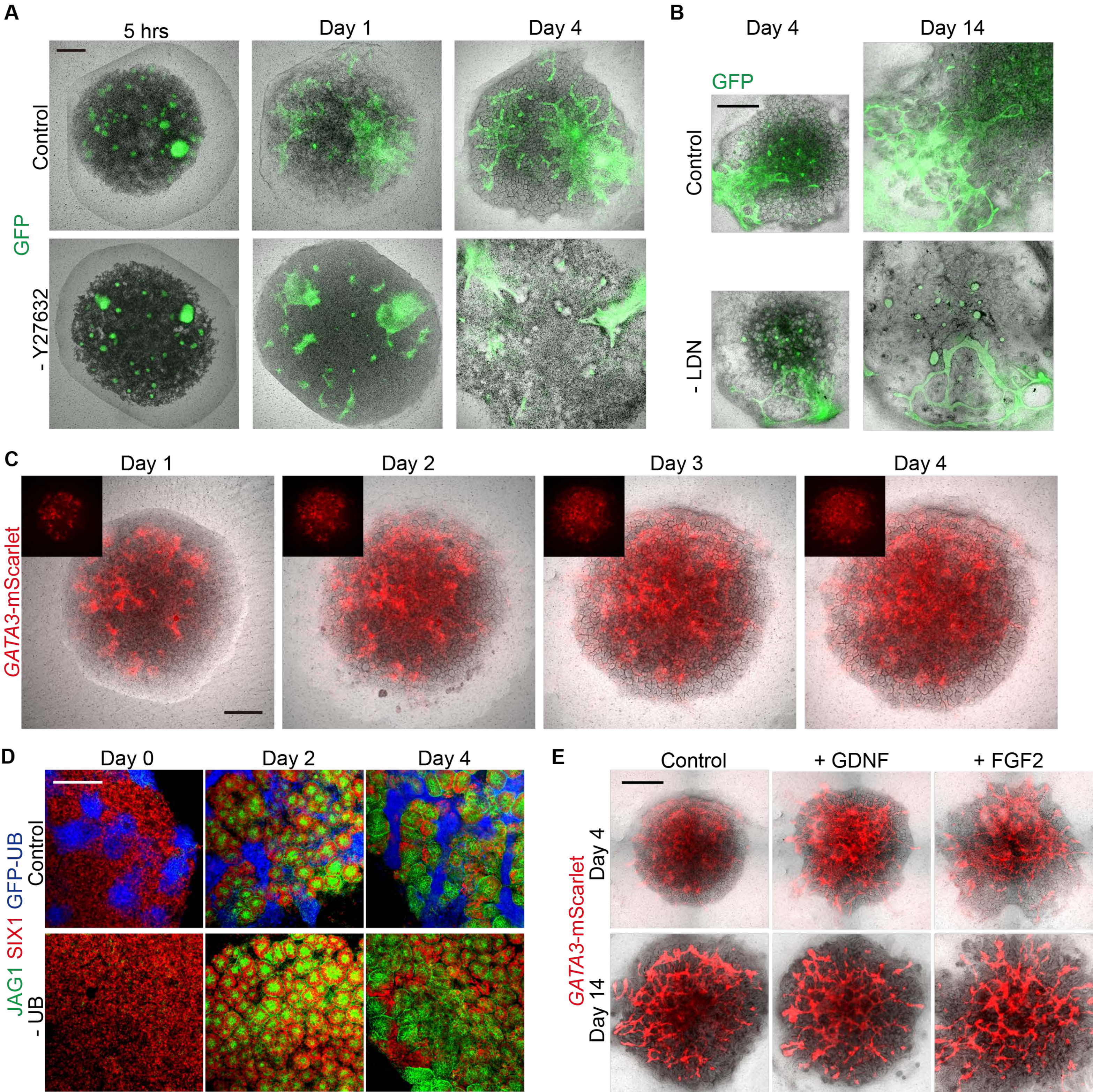

### Supplementary Figure 3

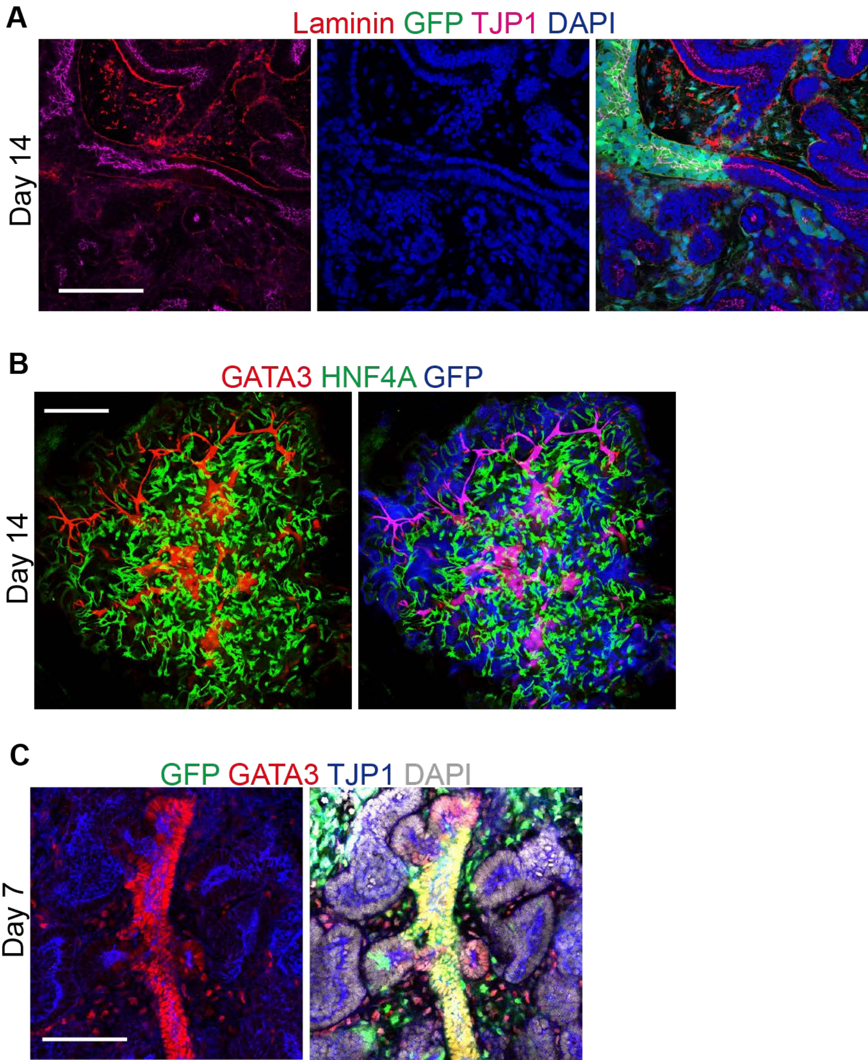

#### Supplementary Figure 4

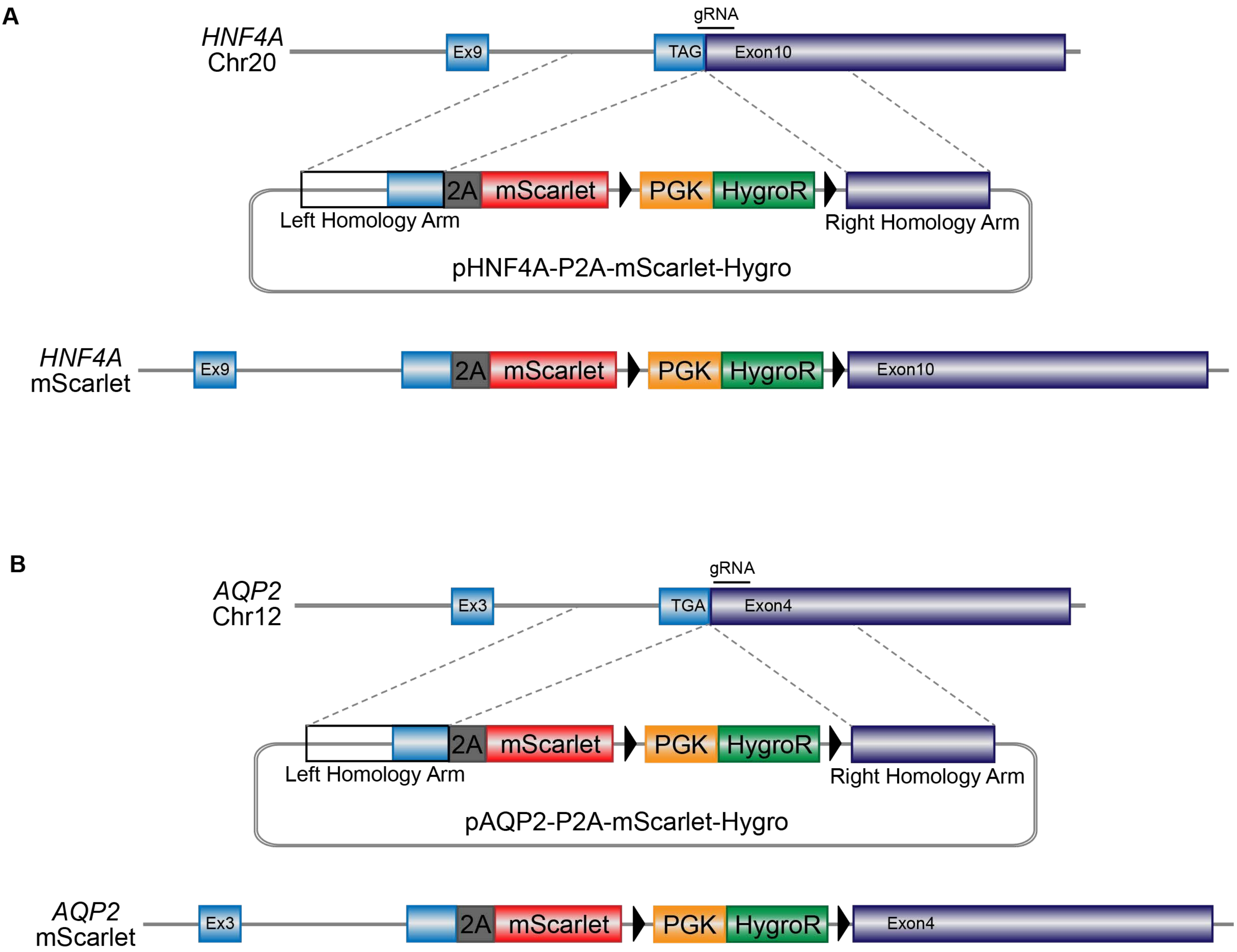

#### Supplementary Figure 5

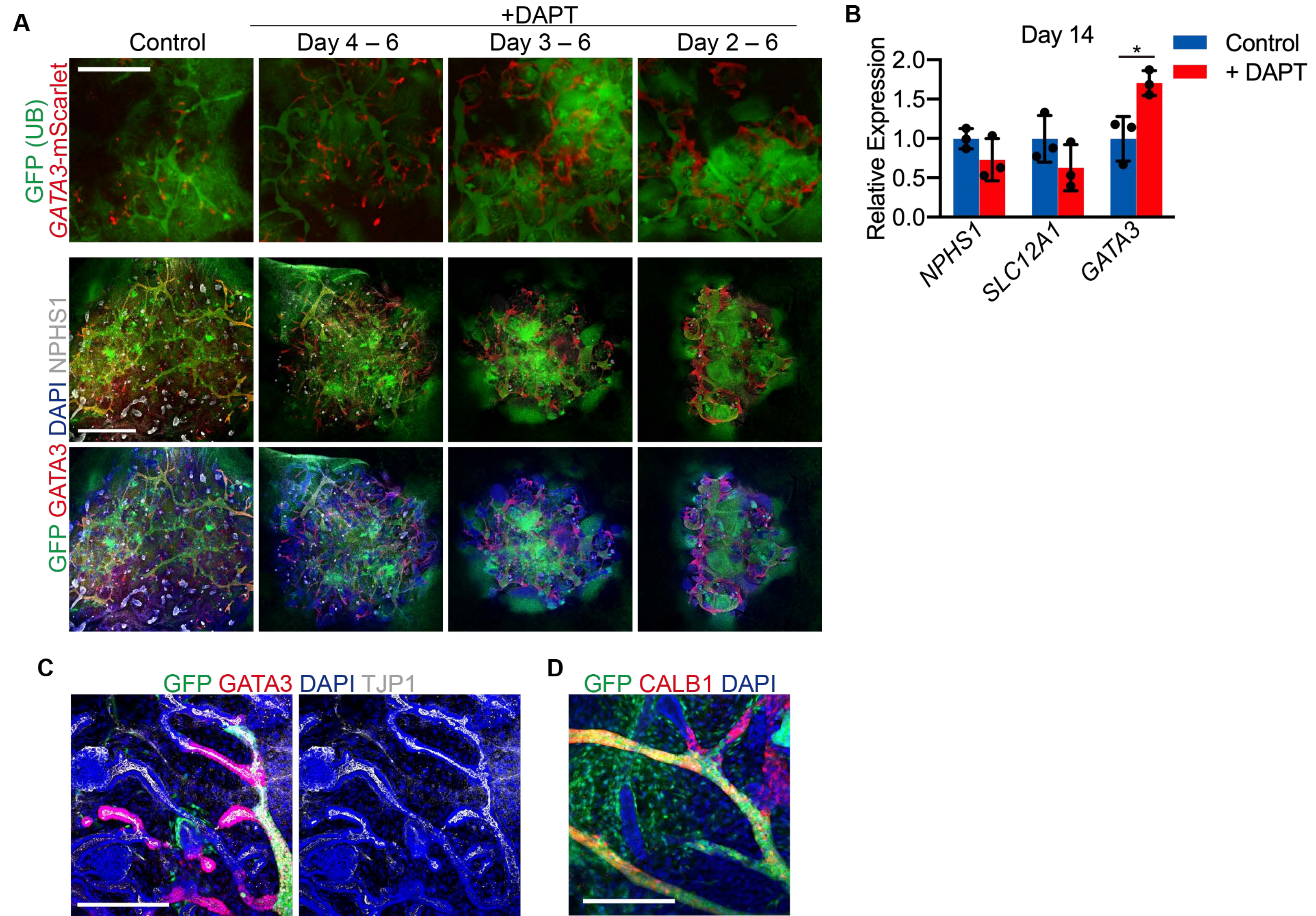

Supplementary Figure 6

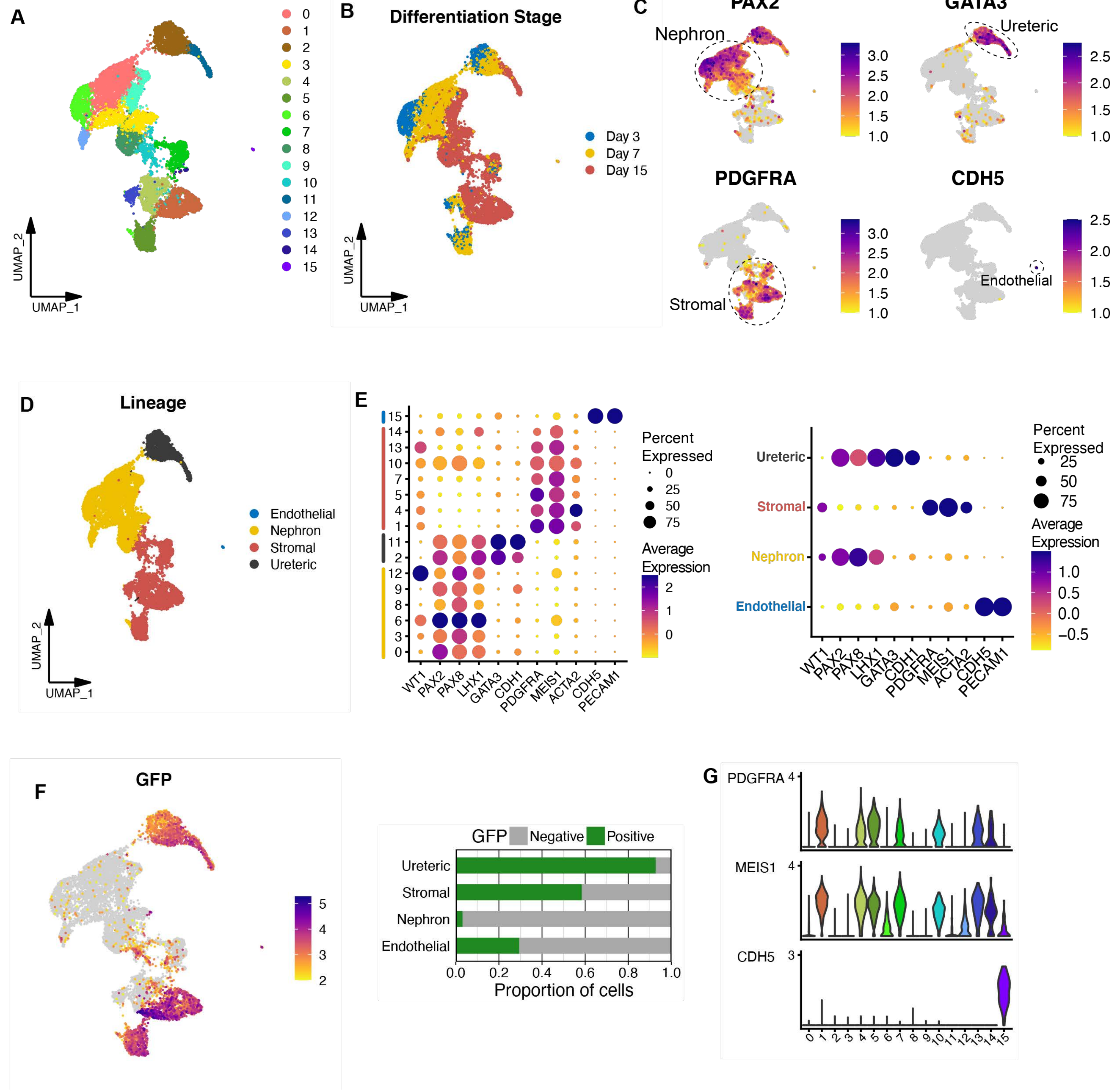

#### Supplementary Figure 7

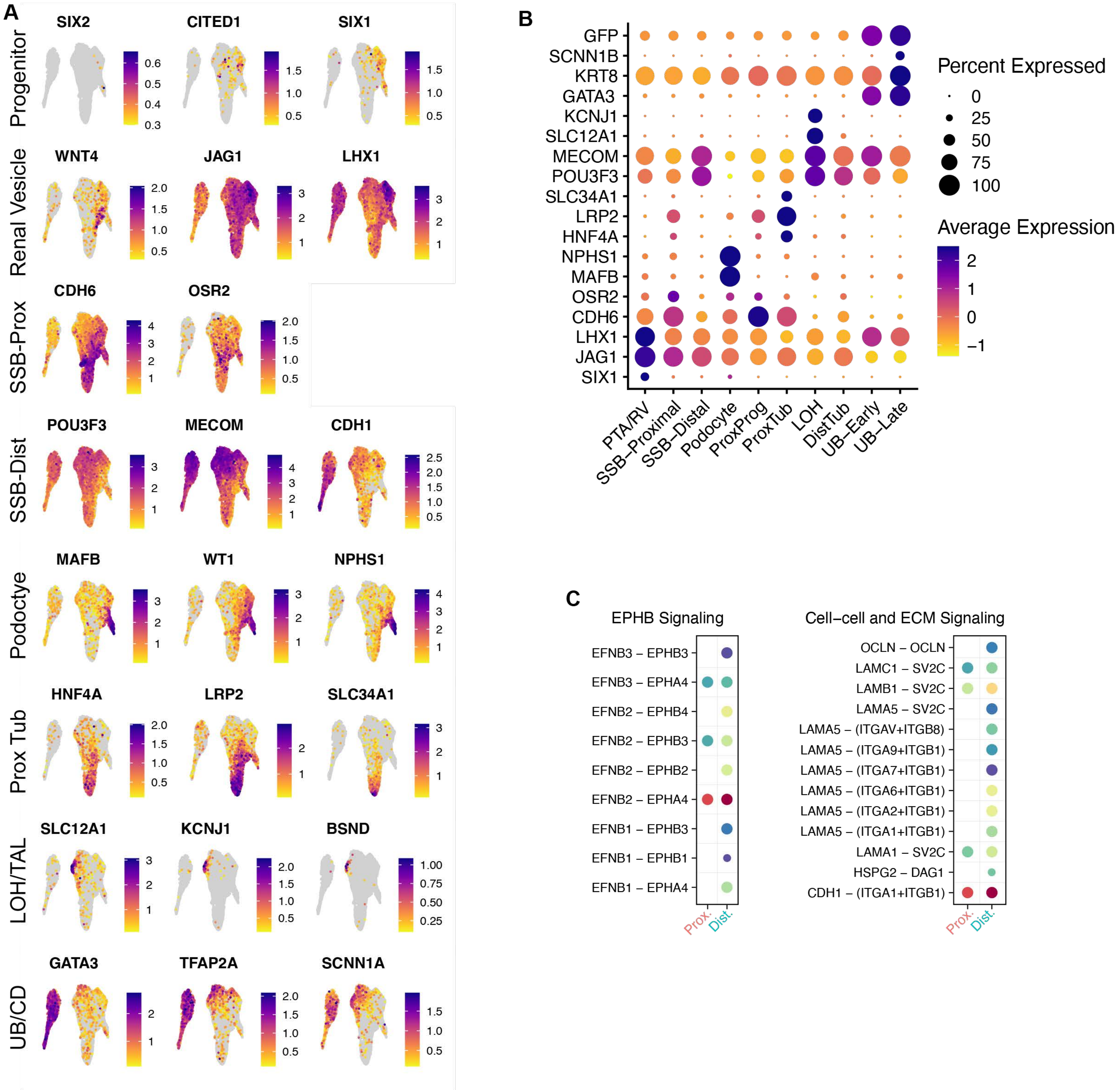

**Supplementary Figure 8**

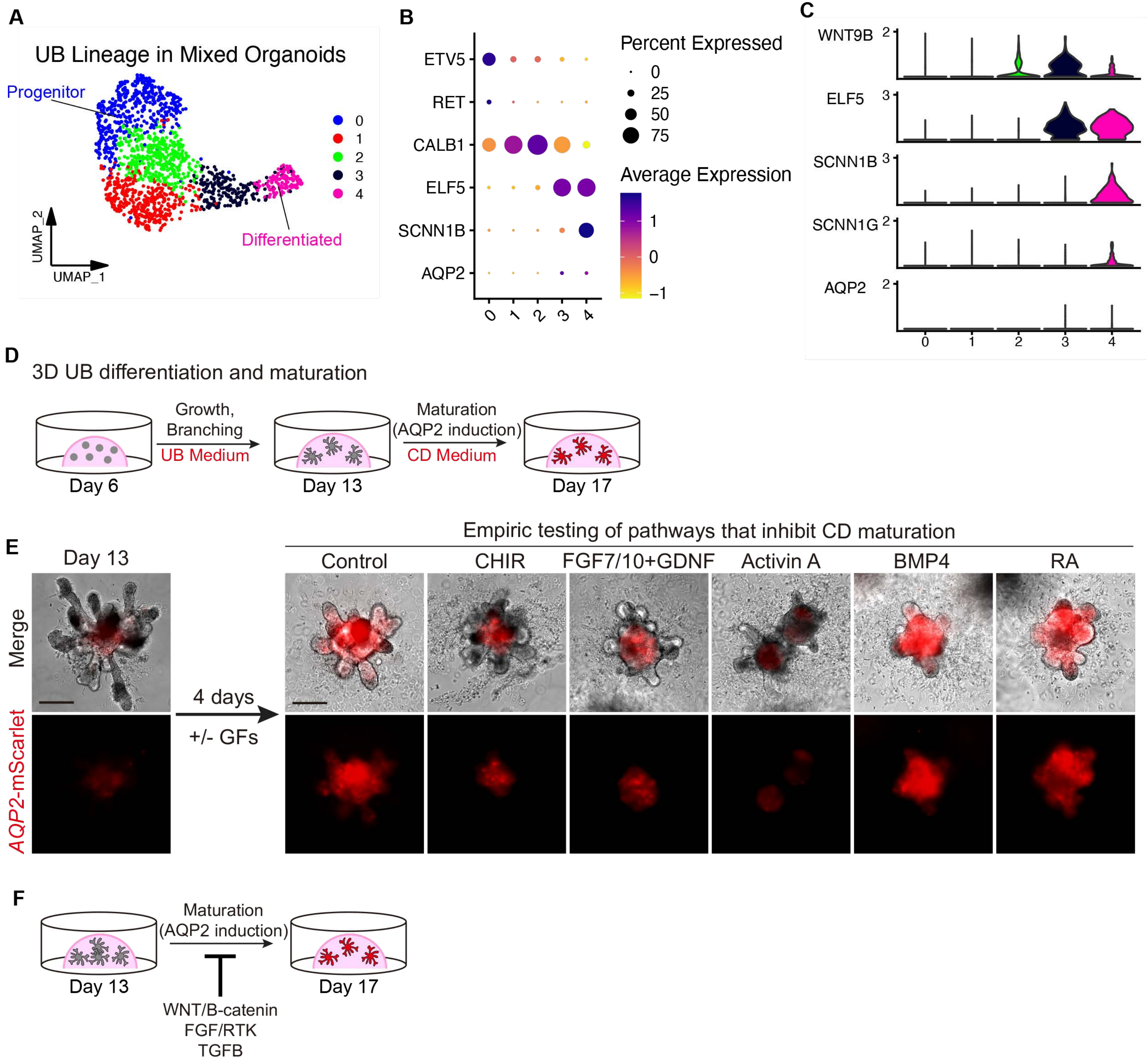

### Supplementary Figure 9

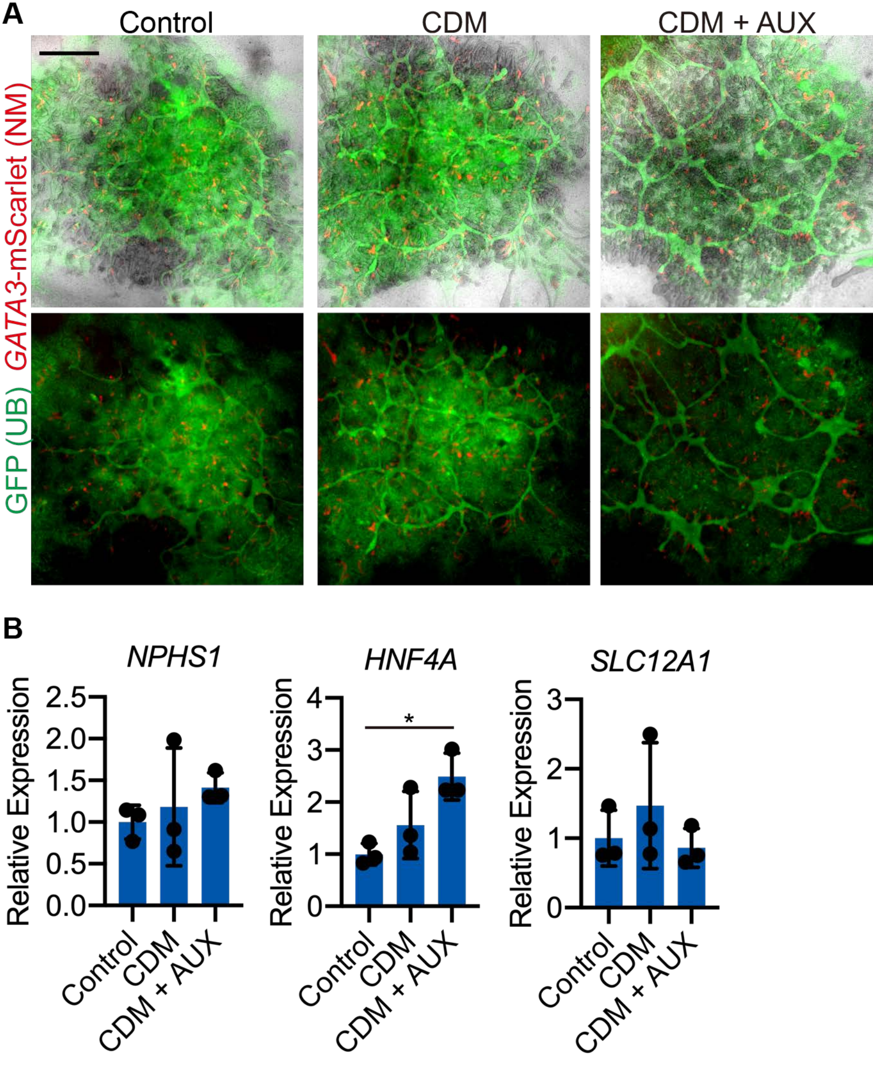
